# The ribosome-associated chaperone Zuo1 controls translation upon TORC1 inhibition

**DOI:** 10.1101/2022.12.09.519716

**Authors:** Ailsa Black, Thomas D Williams, Houjiang Zhou, Adrien Rousseau

## Abstract

Eukaryotic cells achieve proteostasis by ensuring their protein requirements are met through tight control of TORC1 activity. Upon TORC1 inhibition, degradative activity is increased, and protein synthesis is reduced through inhibition of translation initiation, to maintain cell viability. Here, we show that the ribosome-associated complex (RAC)/Ssb chaperone system is required to maintain proteostasis and cell viability under TORC1 inhibition, in yeast. In the absence of the Hsp40 cochaperone Zuo1, translation does not decrease in response to loss of TORC1 activity. The functional interaction between Zuo1 and its Hsp70 partner, Ssb, is required for proper translational control and proteostasis maintenance upon TORC1 inhibition. Further, we have found that the rapid degradation of eIF4G following TORC1 inhibition is prevented in *zuo1Δ* cells, contributing to decreased survival in these conditions. Our findings suggest a new role for RAC/Ssb in regulating translation in response to changes in TORC1 signalling.

## INTRODUCTION

Cells must regularly adapt their proteome in order to respond to changing environmental conditions. The coordination of cellular processes to maintain a stable and functional proteome is known as protein homeostasis (proteostasis), the dysregulation of which has been linked to various disease states, including neurodegenerative disorders (1, 2).

A key regulator of the proteostasis network is the target of rapamycin complex 1 (TORC1), which adapts the equilibrium between catabolic and anabolic processes in response to environmental signals (3). In nutrient rich conditions, TORC1 promotes transcription, translation, and ribosome biogenesis (4–6), while restricting proteasomal and autophagic activity (7–9), to ensure cellular protein requirements for growth are met. Additionally, a network of chaperones ensure that proteins reach their native conformation and prevent the accumulation of protein aggregates (10). Nascent polypeptide chains are particularly vulnerable to misfolding and aggregation. Therefore, eukaryotes utilise a specialised chaperone system composed of the ribosome associated complex (RAC) and the Ssb class of Hsp70 chaperones in yeast (cytosolic Hsp70 in mammalian cells) (11–13). The RAC is anchored near the ribosome exit tunnel and consists of the Hsp40 chaperone, Zuo1 (ZRF1/MPP11 in mammals), and the atypical Hsp70, Ssz1 (HSP70L1 in mammals). The RAC recruits Ssb1 and Ssb2 (collectively known as Ssb) and stimulates their ATPase activity, facilitating their interaction with nascent chains as they emerge from the exit tunnel, protecting them from misfolding and aggregation (14, 15).

In response to stress, the proteome must be rapidly rewired to execute stress survival mechanisms (16–18). The inhibition of TORC1 under various stresses including nutrient deprivation, assists with this essential rewiring and stress response. Protein degradation is increased through upregulation of proteasome activity and autophagy induction, increasing the pool of available amino acids which can be redirected towards the synthesis of stress response proteins (7, 8). Complementing this is a global reduction of translation, which is often accompanied by translational reprogramming, resulting in only selective translation of proteins required for stress survival (9, 19–22). In addition to that, most cytosolic chaperones are transcriptionally upregulated in response to a variety of stresses to enable protein folding to occur under more difficult conditions, with several having a well-documented role in responding to proteotoxic stress (23). Notably, this is not the case for Zuo1 which is coregulated with the translational machinery and has not yet been implicated in the refolding of stress-denatured proteins. Interestingly, it has been found that *zuo1Δ* cells are sensitive to the selective inhibitor of TORC1, rapamycin (22). However, the role of Zuo1 in responding to TORC1 inhibition is unknown.

Here we investigate the importance of Zuo1 for maintaining proteostasis following TORC1 inhibition in S. cerevisiae. We demonstrate a role of Zuo1 in attenuating the rate of translation following TORC1 inhibition. This is dependent on its role as a ribosome associated co-chaperone for Ssb. We further identify Zuo1 interaction partners and found that one candidate, the translation initiation factor eIF4G2 (also known as TIF4632), is misregulated following TORC1 inactivation in *zuo1Δ* cells. This contributes to the loss of cell growth upon TORC1 inhibition. Together, these results define that the RAC-Ssb complex is fine-tuning the rate of translation upon TORC1 inhibition, so that proteostasis, and thereby cell viability, is maintained.

## RESULTS

### Proteostasis is impaired upon TORC1 inhibition in *zuo1Δ* cells

It has previously been shown that loss of Zuo1 sensitizes cells to the TORC1 inhibitor rapamycin (23). In agreement with this, we find that functionality of the RAC is important for surviving rapamycin challenge, as loss of either complex member results in increased sensitivity to rapamycin, with *zuo1Δ* cells displaying a more acute sensitivity than *ssz1Δ* cells (Figure 1A and 1B). As TORC1 is a central coordinator of the protein homeostasis network, we analysed if proteostasis was disrupted by loss of Zuo1, by assessing the load of polyubiquitinated conjugates. A decrease of polyubiquitinated conjugates was seen in WT cells following TORC1 inhibition (Figure 1C), resulting from increased protein degradation and translation inhibition (13, 16). However, this decrease in polyubiquitinated proteins following rapamycin treatment was not observed in *zuo1*Δ cells. This was despite TORC1 being inhibited to a similar extent as in WT cells, with TORC1 activity assessed by phosphorylation of its target Rps6 (Figure 1C), as previously described (24). To examine the clearance of misfolded cytosolic proteins, we utilised the misfolded mutant of carboxypeptidase Y (CPY*) fused to GFP and lacking its endoplasmic reticulum targeting signal (Δss-CPY*GFP) as a reporter (25). This reporter is continually degraded by the proteasome but becomes stabilized in cells with a defect in the ubiquitin proteasome system. In agreement with the impaired clearance of ubiquitinated proteins observed, clearance of Δss-CPY*GFP, is delayed in *zuo1Δ* cells, following rapamycin treatment (Figure 1D and 1E). Together, this indicates that *zuo1Δ* cells are unable to properly adapt their proteostasis network in response to TORC1 inhibition by rapamycin.

**Figure 1.**
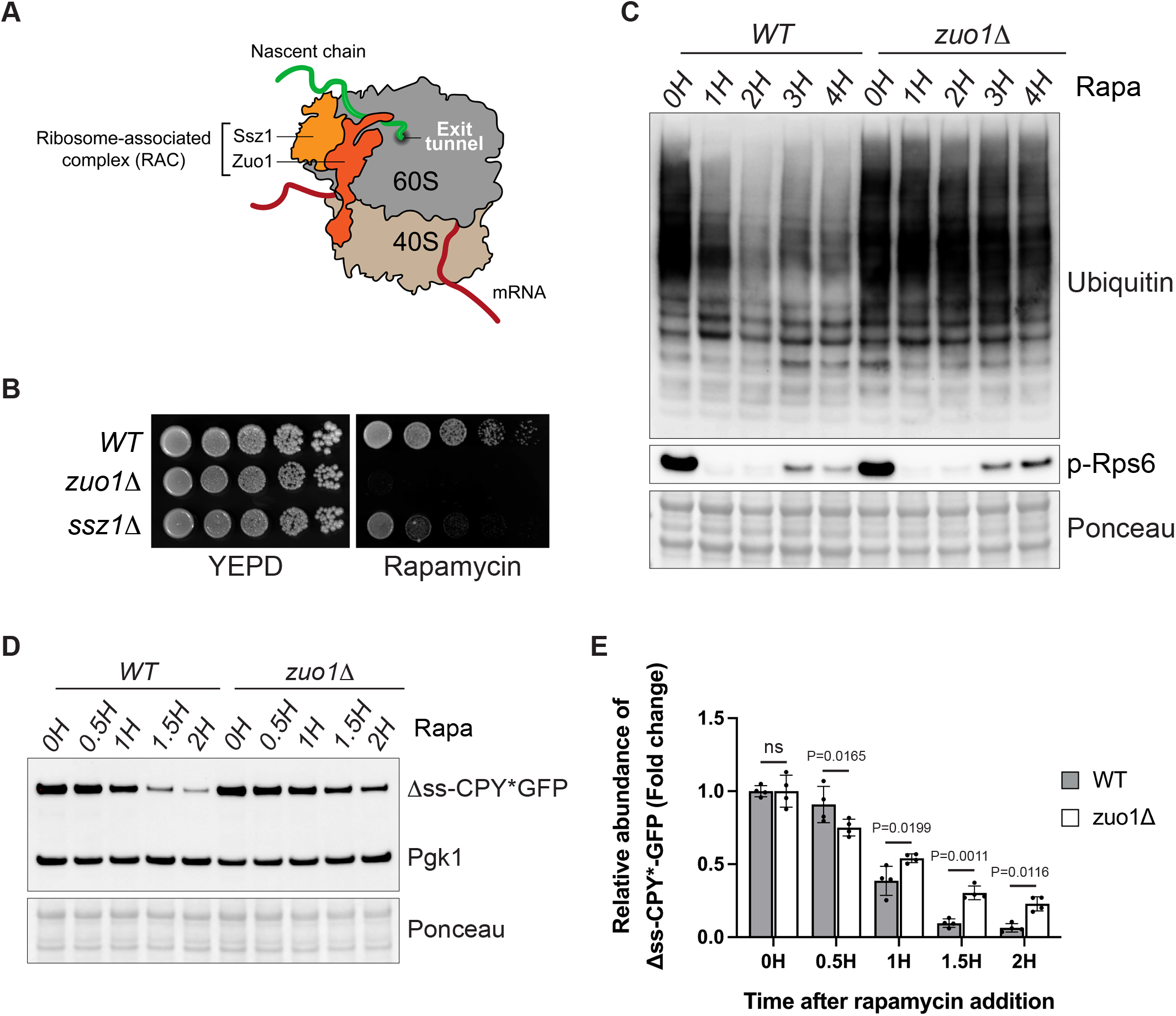
Zuo1 controls protein homeostasis upon TORC1 inhibition. **(A)** Cartoon depicting the Ribosome Associated Complex. Zuo1 interacts with both 40S and 60S ribosomal subunits and Ssz1 remains associated with the ribosome through its interaction with Zuo1. The RAC sits near the polypeptide exit tunnel where it can interact with nascent chains. **(B)** Fivefold serial dilutions of the indicated strains grown on YEPD plates with or without 20ng/ml rapamycin for 4 days at 30°C. **(C)** Immunoblot analysis of lysates from WT and *zuo1*Δ cells treated with 200nM rapamycin for the indicated time. Ponceau staining served as the loading control. **(D)** WT and *zuo1Δ* cells expressing Δss-CPY*GFP from a plasmid were treated with 200nM rapamycin for the indicated time. The resulting lysates were analysed by immunoblotting. Ponceau and Pgk1 staining served as the loading control. **(E)** Graph shows densitometry analysis (mean ± s.d.) of the relative abundance of Δss-CPY*GFP (normalised to Pgk1 levels) from (D) relative to the 0H time point. Statistical significance was assessed using two-way ANOVA t-test (n = 4 independent biological replicates). n. s. (not significant).

### Proteasomal degradation is unaffected in *zuo1Δ* cells

Impaired clearance of polyubiquitinated proteins can be a sign of defective protein degradation by the proteasome. Following TORC1 inhibition, cells boost their degradative capacity by increasing proteasome assembly and activity (8, 26). To assess proteasome function in *zuo1Δ* cells, we performed an in-gel peptidase assay. The advantage of this method is that it can resolve the activity of all proteasomal species separated on a native gel after adding an internally quenched fluorogenic peptide. Fluorescence from the peptide is only observed after cleavage by the proteasome, and hence fluorescence intensity mirrors proteasome activity. The increase in proteasome activity following rapamycin treatment is not compromised in *zuo1Δ* cells, and this was true for both singly-and doubly-capped proteasomes (RP-CP and RP_2_-CP, respectively) (Figure 2A). In agreement, when ribosomal translation is inhibited by cycloheximide the degradation of the proteasome reporter substrate ΔssCPY*GFP is comparable in WT and *zuo1Δ* cells (Figure 2B and 2C). Thus, the loss of proteostasis in *zuo1Δ* cells is not due to a defect in proteasomal degradation.

**Figure 2.**
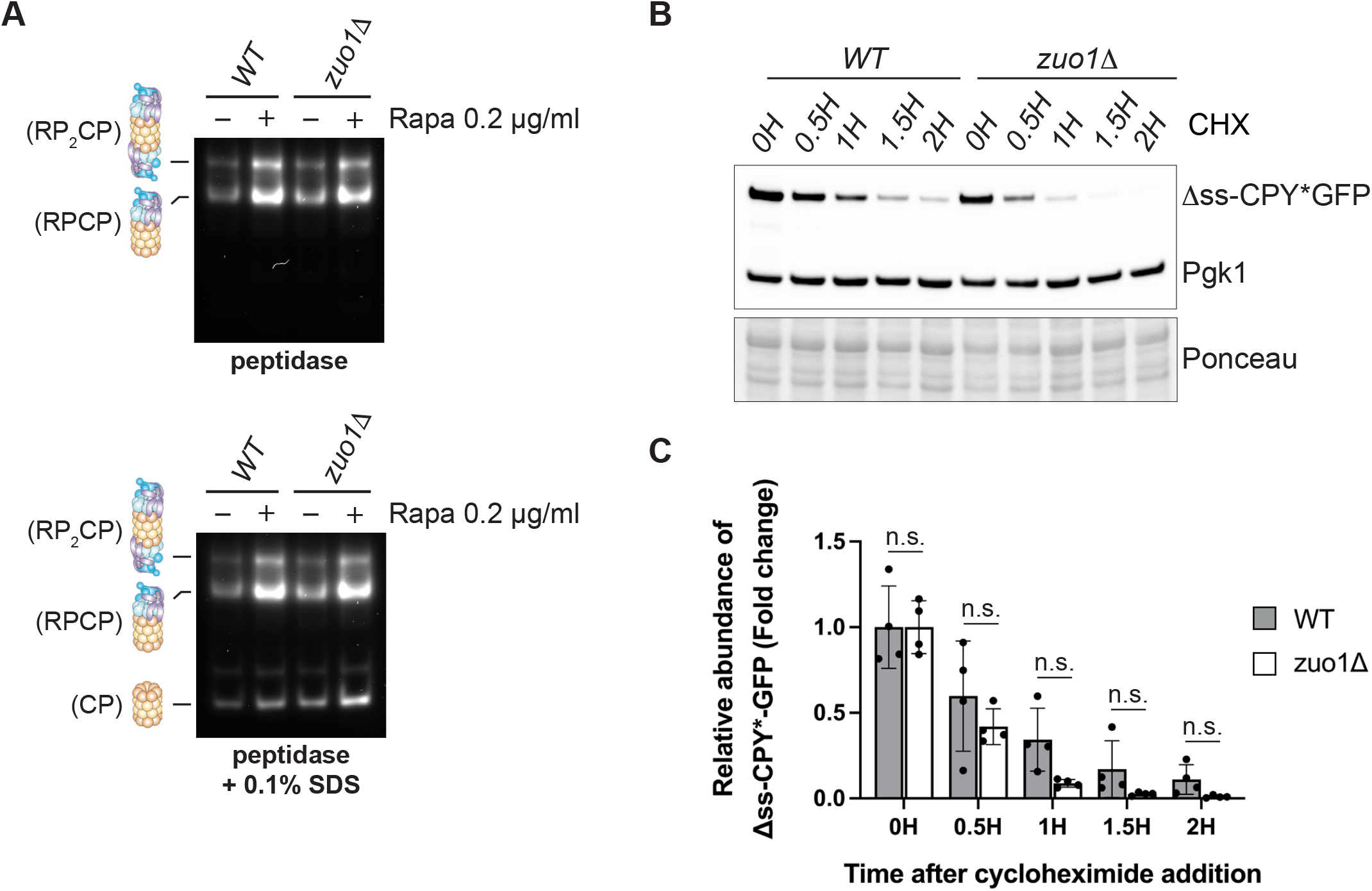
Proteasome homeostasis is not impaired in *zuo1*Δ cells. **(A)** Yeast extracts from cells treated with 200nM rapamycin for 3h or left untreated were separated by native-PAGE (3.8-5% gradient) and peptidase activity detected using the fluorogenic substrate Suc-LLVY-AMC in the presence or absence of 0.1% SDS. RP_2_CP, double-capped proteasome; RPCP, single-capped proteasome; and CP, core particle complexes are indicated. **(B)** Immunoblot analysis of lysates from WT and *zuo1*Δ cells expressing Δss-CPY*GFP from a plasmid treated with 35μg/ml cycloheximide for the indicated time. Ponceau and Pgk1 staining served as the loading control. **(C)** Graph shows densitometry analysis (mean ± s.d.) of the relative abundance of Δss-CPY*GFP (normalised to Pgk1 levels) from (D) relative to the 0H time point. Statistical significance was assessed using two-way ANOVA t-test (n = 4 independent biological replicates). n. s. (not significant).

### Translation is not reduced in *zuo1Δ* mutants following TORC1 inhibition

A possible explanation as to why Δss-CPY*GFP levels remain higher in rapamycin-treated *zuo1*Δ cells, compared to WT cells, despite the clearance of this reporter being unimpeded, is that translation may be continuing despite TORC1 inhibition. Failure to reduce translation following rapamycin treatment would lead to excess production of unwanted proteins, overwhelming the proteostasis network. To test this hypothesis, we employed the SUnSET assay in which the aminoacyl tRNA analog, puromycin, is incorporated into nascent chains, reflecting the rate of translation *in vivo* (27). Following rapamycin treatment, translation decreases in WT cells, correlating with TORC1 inhibition (Figure 3A). Strikingly, rapamycin treatment fails to decrease translation in *zuo1Δ* cells following rapamycin-induced TORC1 inhibition. Correspondingly, a decrease in polysome levels is seen in WT cells following rapamycin treatment (Figure S1A). While *zuo1Δ* cells have lower polysome levels in basal conditions (27, 28), this does not seem to affect the *in vivo* translation rate (Figure 3A). This is consistent with previous studies reporting that overall translation rate is not impaired in RAC-deleted cells, despite polysome levels being lower (28, 29). Furthermore, the decrease in polysome levels observed in WT cells upon TORC1 inhibition, is not present in *zuo1Δ* cells (Figure S1B), suggesting that a failure to properly regulate translation may comprise the proteostasis defect detected. Confirming this, cycloheximide treatment inhibited translation in both WT and *zuo1Δ* cells, even in the presence of rapamycin (Figure 3B), and rescued the impaired clearance of polyubiquitinated proteins observed in *zuo1*Δ cells following TORC1 inhibition (Figure 3C). Similarly, while clearance of Δss-CPY*GFP is impeded in *zuo1Δ* cells treated with rapamycin alone, cotreatment with cycloheximide restores the clearance of the misfolded proteasome reporter in *zuo1*Δ cells following TORC1 inhibition (Figure 1D, 1E, 3D and 3E). Together, this data demonstrates that the failure to shut down translation in response to TORC1 inhibition likely underpins the protein homeostasis defect in *zuo1Δ* cells

**Figure 3.**
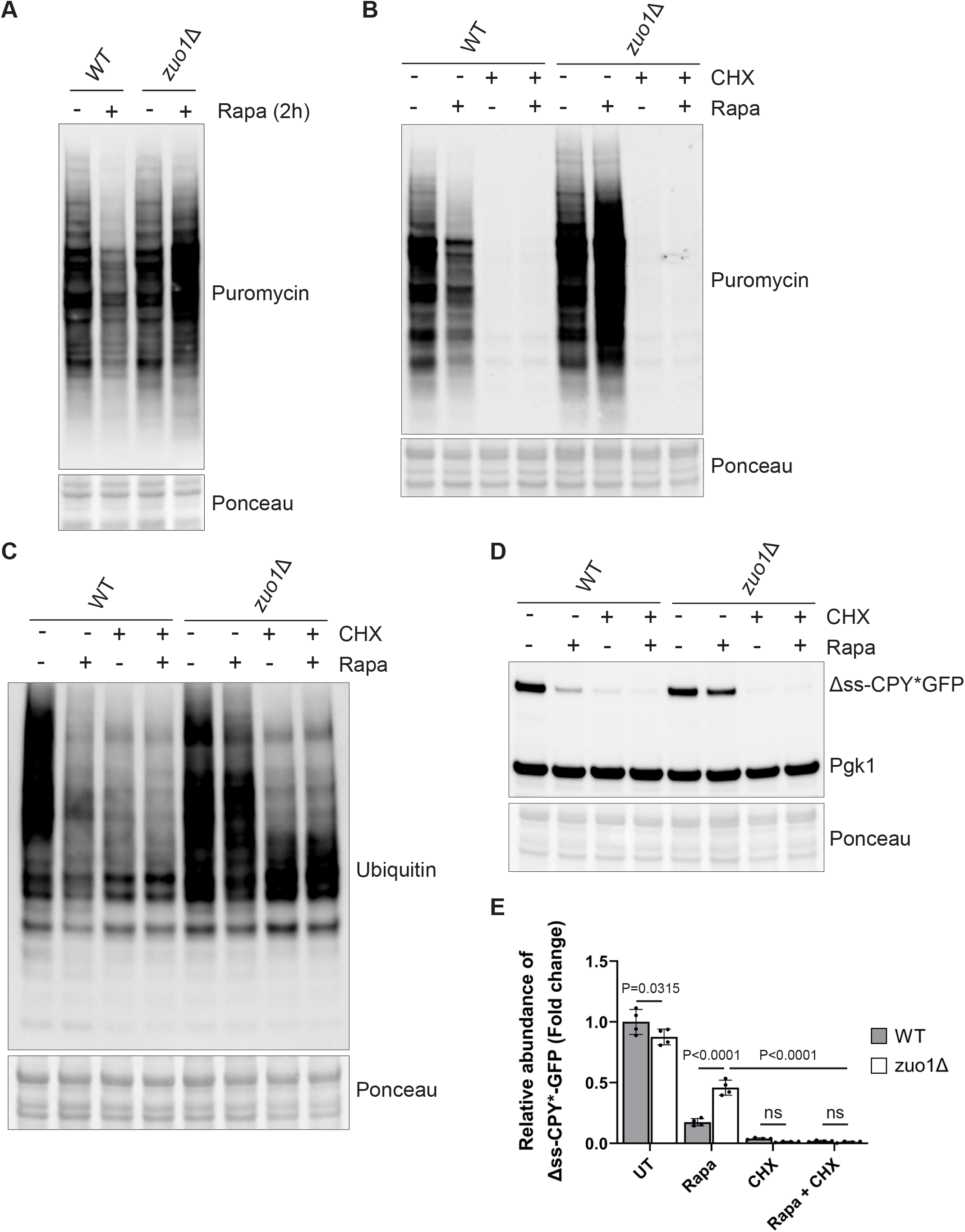
Translation is not reduced in *zuo1*Δ cells upon TORC1 inhibition. **(A)** Immunoblots of lysates from WT and zuo1Δ cells grown in the presence of 0.5mM puromycin, with or without 200nM rapamycin for 2 hours. Ponceau staining served as the loading control. **(B)** Immunoblot analysis of lysates from WT and *zuo1*Δ cells grown in the presence 0.5mM puromycin and treated with 200nM rapamycin and/or 35μg/ml cycloheximide as indicated, for 2 hours. Ponceau staining served as the loading control. **(C)** Immunoblot analysis of lysates from WT and *zuo1*Δ cells treated with 200nM rapamycin and/or 35μg/ml cycloheximide as indicated, for 2 hours. Ponceau staining served as the loading control. **(D)** Immunoblot analysis of lysates from WT and *zuo1*Δ cells expressing Δss-CPY*GFP from a plasmid treated with 200nM rapamycin and/or 35μg/ml cycloheximide for the indicated time. Ponceau and Pgk1 staining served as the loading control. **(E)** Graph shows densitometry analysis (mean ± s.d.) of the relative abundance of Δss-CPY*GFP (normalised to Pgk1 levels) from WT and *zuo1*Δ cells treated with 200nM rapamycin and/or 35μg/ml cycloheximide as indicated, for 2 hours. Statistical significance was assessed using two-way ANOVA t-test (n = 4 independent biological replicates). n. s. (not significant).

### Zuo1 controls translation upon TORC1 inhibition through Ssb chaperones

Having established the importance of Zuo1 in mediating a translational decrease following TORC1 inhibition, we set out to define how it achieves this. Zuo1, as a key component of the RAC, functions as a cochaperone for Ssb. However, Zuo1 is also reported to have additional functions (Figure 4A). The first is in recruitment of Ltn1, a key E3 ligase of ribosome associated quality control (30), and the second is in the induction of pleiotropic drug resistance, through the Pdr1 transcription factor (31). Deletion of Ltn1 or Pdr1 does not confer sensitivity to rapamycin (Figure 4B) or result in impaired clearance of polyubiquitinated proteins (Figure 4C), suggesting that these roles of Zuo1 are dispensable for its role in adapting the proteostasis network in response to TORC1 inhibition. Deletion of both homologs of Ssb (ssb1 and ssb2), however, results in acute sensitivity to rapamycin and is accompanied by a defect in clearing polyubiquitinated proteins following TORC1 inhibition, analogous to *zuo1Δ* cells (Figure 4B and 4C). Cells lacking Ssb also fail to shut down translation in response to TORC1 inhibition (Figure 4D). As in *zuo1Δ* cells, this seems to underpin the proteostasis defect, as restoring translation inhibition with cycloheximide in *ssb1/2Δ* cells improves their clearance of polyubiquitinated proteins upon rapamycin treatment (Figure 4E).

**Figure 4.**
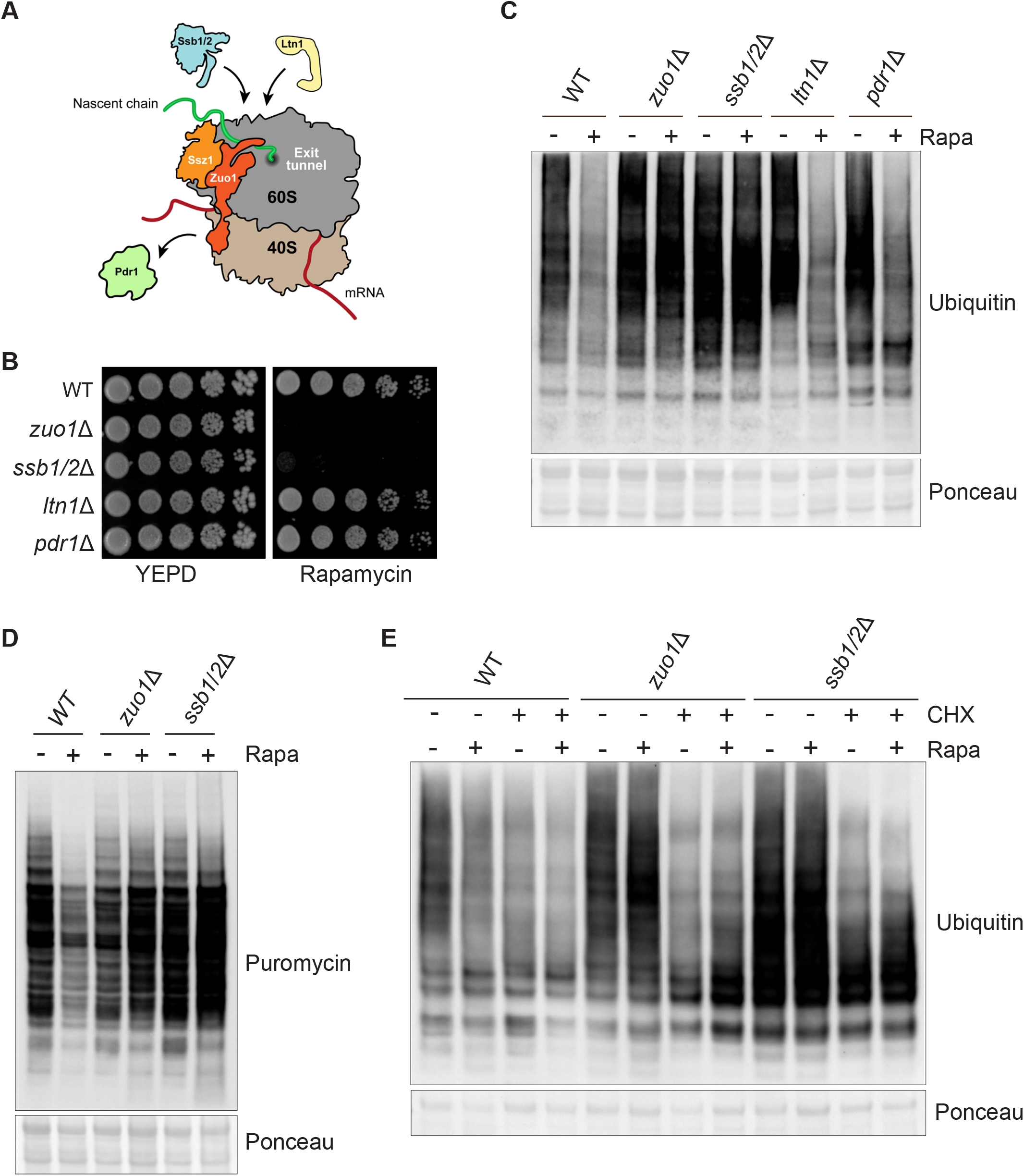
Ssb reduces translation upon TORC1 inhibition. **(A)** Cartoon illustrating the known roles of Zuo1. Zuo1 is the cochaperone of Ssb, recruiting it to the ribosome and stimulating its ATPase activity. The recruitment of a key ubiquitin ligase in ribosome-associated quality control, Ltn1, to the ribosome is facilitated by Zuo1. The transcription factor Pdr1 can be activated by Zuo1 when it is not ribosome associated and the C-terminal 4-helix bundle is unfolded. **(B)** Cells of the indicated strains were spotted in five-fold serial dilutions onto YEPD plate +/-20ng/ml rapamycin and grown for 4 days at 30°C. **(C)** Immunoblot analysis of lysates from cells of the indicated genotype treated with 200nM rapamycin for 4 hours or left untreated. Ponceau staining served as the loading control. **(D)** Immunoblot analysis of lysates from WT, *zuo1*Δ, and *ssb1/2*Δ cells grown in the presence of 0.5mM puromycin, with or without 200nM rapamycin for 2 hours. Ponceau staining served as the loading control. **(E)** Immunoblot analysis of lysates from cells of the indicated genotype treated with 200nM rapamycin and/or 35μg/ml cycloheximide as indicated, for 2 hours. Ponceau staining served as the loading control.

*zuo1Δ* and *ssb1/2Δ* cells share sensitivity to many of the same stresses and drugs, such as translation inhibitors and cold stress (32). For many of these, it has been demonstrated that the sensitivity of *zuo1Δssb1/2Δ* cells is the same as that of *zuo1Δ* cells and is not additive (32). Likewise, we have found that the phenotype of *zuo1Δssb1/2Δ* cells resembles that of cells lacking either Zuo1 or Ssb, displaying similarly low levels of growth and impaired polyubiquitin clearance upon rapamycin treatment (Figures 5A and 5B). Additionally, translation is not shut down in *zuo1Δssb1/2Δ* cells following TORC1 inhibition, but the overall phenotype is not more severe than either *zuo1*Δ cells or *ssb1/2Δ* cells (Figure 5C). This further emphasises that the role of Zuo1 in maintaining proteostasis upon TORC1 inhibition is dependent on its role as a ribosome associated co-chaperone.

**Figure 5.**
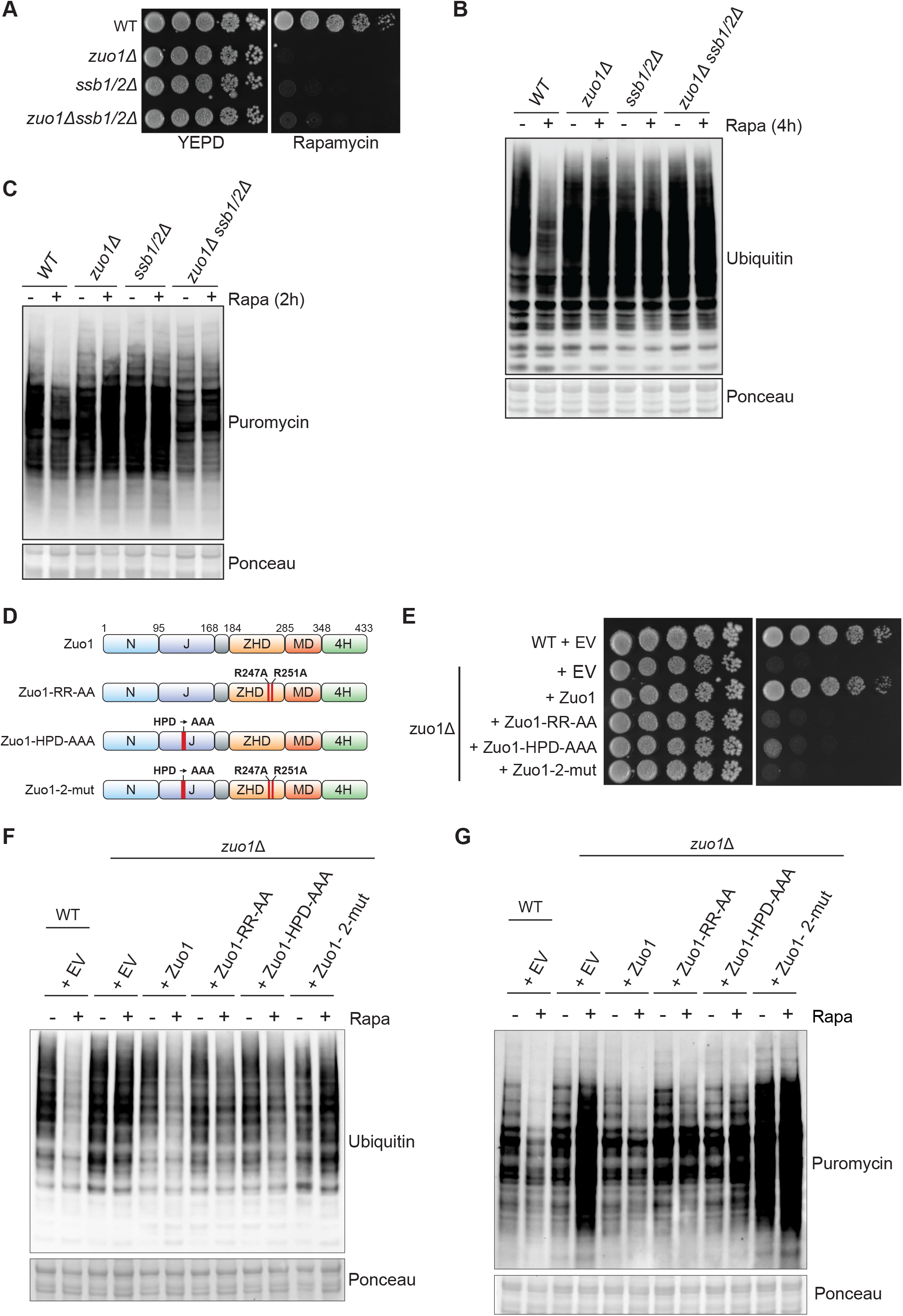
Zuo1-mediated Ssb regulation controls translation upon TORC1 inhibition. **(A)** Cells of the indicated strains were spotted in five-fold serial dilutions onto YEPD plate +/-20ng/ml rapamycin and grown for 4 days at 30°C. **(B)** Immunoblot analysis of lysates from cells of the indicated genotype treated with 200nM rapamycin for 4 hours or left untreated. Ponceau staining served as the loading control. **(C)** Immunoblot analysis of lysates from the indicated strains grown in the presence of 0.5mM puromycin, with or without 200nM rapamycin for 2 hours. Ponceau staining served as the loading control. **(D)** Zuo1 mutants used in this study. The domain architecture of Zuo1 is illustrated with mutated residues highlighted in red. N-terminal domain (N), J-domain (J), Zuotin homology domain (ZHD), middle domain (MD), four-helix bundle (4H). **(E)** WT and *zuo1*Δ cells expressing the indicated plasmids or empty vector (EV) were spotted in five-fold serial dilutions onto YEPD plate +/-20ng/ml rapamycin and grown for 4 days at 30°C. **(F)** Immunoblot analysis of lysates from WT and *zuo1*Δ cells expressing the indicated plasmids with or without 200nM rapamycin for 2 hours. Ponceau staining served as the loading control. **(G)** Immunoblot analysis of lysates from WT and *zuo1*Δ cells expressing the indicated plasmids or empty vector (EV) were grown in the presence of 0.5mM puromycin, with or without 200nM rapamycin for 2 hours. Ponceau staining served as the loading control.

To confirm this result, we utilised two mutations in Zuo1 which ablate crucial functions in regulating Ssb: a ribosome binding mutant in which two key arginine residues (R247/R251) are mutated to alanine, destabilizing the interaction of Zuo1 with ribosomes (RR-AA) (33), and a HPD-AAA mutant in which the conserved HPD motif in the J domain of Zuo1 has been mutated, inhibiting its ability to stimulate the ATPase activity of Ssb (Figure 5D) (34, 35). Expression of WT Zuo1 on a plasmid complemented the growth defect of *zuo1Δ* cells in response to rapamycin, whereas expression of either of the RR-AA or HPD-AAA Zuo1 mutants did not (Figure 5E). However, the rapamycin sensitivity of *zuo1Δ* cells re-expressing either of these Zuo1 mutants is not as severe as cells expressing the control plasmid (EV: Empty Vector). Similarly, relative to WT cells, the clearance of ubiquitinated proteins is impaired in *zuo1Δ* cells expressing either the ribosome binding or chaperone mutant of Zuo1, but to a lesser extent than in *zuo1Δ* cells harbouring the control plasmid (Figure 5F). Translation is also not fully shut down following TORC1 inhibition in cells expressing the single Zuo1 mutants, however, the phenotype is not as severe as that displayed by *zuo1Δ* cells (Figure 5G). It has been reported that even very low levels of Zuo1 are enough to preserve its function suggesting that single mutations may not be enough to fully abrogate Zuo1 activity (32). Therefore, we generated a double mutant (RR-AA/HPD-AAA; zuo1-2-mut) lacking both ribosome binding and a functional J domain to account for any residual function of the individual mutants (Zuo1-2mut). Cells expressing Zuo1-2mut displayed greater sensitivity to rapamycin than those expressing the single mutants, more akin to deletion of the protein (Figure 5E). Accordingly, expression of Zuo1-2mut was not able to rescue the defective clearance of polyubiquitinated proteins and shutdown of translation observed in *zuo1Δ* cells upon TORC1 inhibition (Figure 5F and 5G). This confirms the importance of Zuo1 function in regulating Ssb chaperones for maintaining proteostasis in response to TORC1 inhibition.

### Identification of Zuo1 partners upon TORC1 inhibition

We have determined that of the known roles of Zuo1, only its role in activating Ssb at the ribosome, appears to be necessary for regulating proteostasis in cells challenged with rapamycin. However, it is unclear if additional Zuo1 partners are important for this regulation. We therefore immunoprecipitated endogenously tagged Zuo1 from cells grown in the presence or absence of rapamycin (Figure 6A). We then performed tandem mass tag (TMT)-based quantitative proteomics to identify novel interactors and analyse changes in its interactions (Figure 6B). All the members of the RAC/Ssb chaperone system and many ribosomal proteins were found to be interacting with Zuo1 in our dataset (all proteins with at least 50% peptide coverage; Figure S2A). A subset of glycolytic enzymes was also identified to be interacting with Zuo1. These genes are highly expressed and, with one exception, have been shown to be RAC dependent substrates of Ssb (15), which likely accounts for their identification as Zuo1 interactors. Furthermore, gene ontology analysis revealed a strong enrichment of proteins linked to translation fidelity as well as those involved in ribonucleoprotein biogenesis and localization, processes which Zuo1 is known to be involved in (Figure S2B). This highlights the robustness of this dataset and gives confidence in the novel Zuo1 interactors identified.

**Figure 6.**
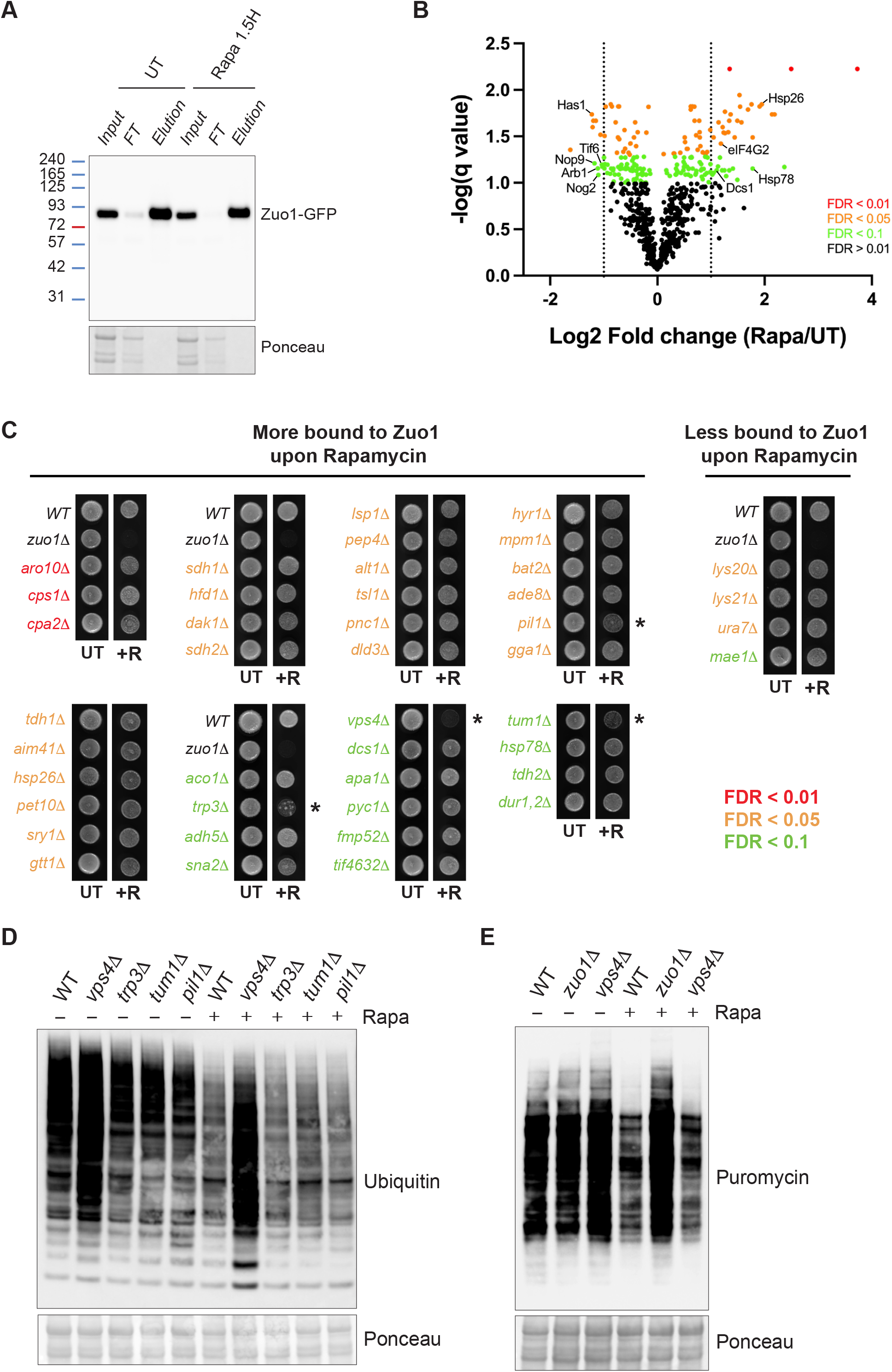
Identification of Zuo1 partners upon TORC1 inhibition. **(A)** Cells expressing endogenously tagged Zuo1-GFP grown in the presence or absence of 200nM rapamycin for 1.5h, subjected to immunoprecipitation using an anti-GFP nanobody and analysed by immunoblotting. **(B)** Volcano plot illustrating Zuo1 interactors from Zuo1-GFP immunoprecipitates, differentially enriched in UT and rapamycin-treated conditions. Red, orange, and green dots represent proteins with a false discovery rate of <0.01, <0.05, and <0.1, respectively. The dotted lines indicate the threshold for two-fold differential enrichment between UT and rapamycin-treated conditions. **(C)** Growth assays of deletion mutants of non-essential Zuo1 interactors from (B) with greater than two-fold change in enrichment between rapamycin and UT conditions. The indicated strains were grown on YEPD plates with or without 20ng/ml rapamycin for 4 days. Strains which display increased sensitivity towards rapamycin compared to WT cells are highlighted with an asterisk (*). **(D)** WT cells and rapamycin sensitive deletion mutants from (C) were treated with 200nM rapamycin for 4 hours or left untreated. The resulting lysates were analysed by immunoblotting. Ponceau staining served as the loading control. **(E)** Immunoblots of lysates from WT, *zuo1*Δ, and *vps4*Δ cells grown in the presence of 0.5mM puromycin, with or without 200nM rapamycin for 2 hours. Ponceau staining served as the loading control.

We identified 39 and 11 proteins whose interaction with Zuo1 was significantly increased and decreased upon rapamycin treatment, respectively. Those which are known to be linked to either translation or chaperone activity have been highlighted in Figure 6B as they are of particular interest for Zuo1’s role in proteostasis. However, in order to prevent bias, we have not limited our search to these proteins. Of the interactions which are significantly changed following rapamycin treatment, we focused on the 43 interactors which are not essential. If these interactors are important for mediating the role of Zuo1 in proteostasis, then it would be expected that their loss would confer a similar phenotype to that of *zuo1Δ* cells upon rapamycin treatment. Deletion mutants of the 43 interactors were initially screened for rapamycin sensitivity, with only *trp3*Δ, *vps4*Δ, *pil1*Δ, and *tum1*Δ cells displaying increased rapamycin sensitivity relative to WT cells (Figure 6C). Subsequently, a proteostasis defect in response to rapamycin challenge was observed in cells lacking Vps4 but not in *trp3*Δ, *pil1*Δ, or *tum1*Δ cells (Figure 6D). However, in *vps4*Δ cells, this defect is not at the level of translation regulation as following TORC1 inhibition translation is still inhibited (Figure 6E). None of the Zuo1 interactors examined seem to be involved in its role in regulating proteostasis following rapamycin treatment. Although the proteostasis defect in *vps4Δ* doesn’t appear to be linked to Zuo1, further study would be valuable to elucidate the defect in the proteostasis network of these cells.

### Degradation of eIF4G is impeded in *zuo1Δ* cells

None of the novel interactors identified appear to be involved in mediating Zuo1’s role in maintaining proteostasis in response to TORC1 inhibition. However, one interactor stood out as being particularly interesting, eIF4G2 (encoded by TIF4632 in yeast), despite the deletion mutant displaying no sensitivity towards rapamycin (Figure 6C). This could be due to the presence of its paralog, eIF4G1 (encoded by TIF4631), as they are functionally redundant. eIF4G1/2 are scaffold proteins in the eIF4F complex which is required for recruiting the 40S ribosomal subunit to mRNA during translation initiation (reviewed in (36)). Translation initiation is the predominant target of TORC1 translational regulation. TORC1 elicits control over this process by promoting the maintenance of eIF4G levels in nutrient rich conditions, facilitating translation initiation. Following TORC1 inhibition, e.g. by rapamycin treatment, eIF4G is rapidly degraded to suppress bulk translation (37). In *zuo1Δ* cells, the dramatic reduction of eIF4G1 and eIF4G2 that is observed in WT cells following rapamycin treatment, is severely impaired (Figure 7A-7D). The protein level of other eIF4F complex members, in both basal and rapamycin treated conditions, is unaffected by loss of Zuo1, indicating that this misregulation is specific to eIF4G (Figure S3). Interestingly, it has been reported that eIF4G is the least abundant of the eIF4 factors and hence is rate-limiting for eIF4F activity (38, 39). This suggests that *zuo1Δ* cells may be sustaining translation through the excess pool of eIF4G.

**Figure 7.**
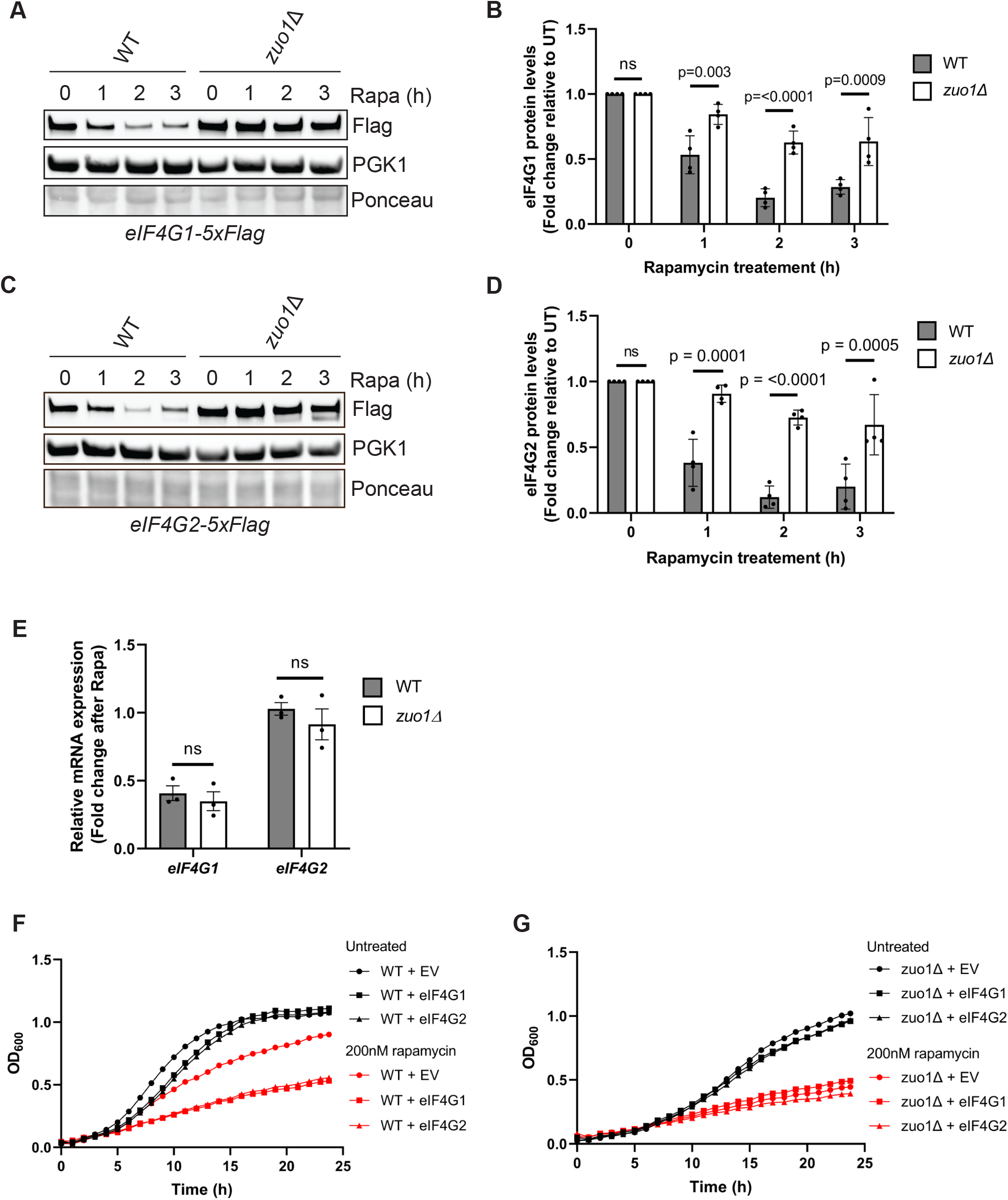
Zuo1 controls eIF4G protein turnover upon TORC1 inhibition. **(A) and (C)** Immunoblot analysis of lysates from WT and *zuo1*Δ cells containing eIF4G1-5xFLAG (**A**) or eIF4G2-5xFLAG (**C**) at the endogenous locus treated with 200nM rapamycin for the indicated time. Ponceau and Pgk1 staining served as the loading control. **(B) and (D)** Graph shows densitometry analysis (mean ± s.d.) of the relative abundance of eIF4G1-5xFLAG and eIF4G2-5xFLAG from (**A**) and (**C**) respectively, relative to the 0h time point. Statistical significance was assessed using two-way ANOVA t-test (n = 4 independent biological replicates). n. s. (not significant). **(E)** mRNA levels of eIF4G1 and eIF4G2 from WT and *zuo1*Δ cells treated with 200nM rapamycin for 2 hours or left untreated were analysed by qRT-PCR. The fold change in mRNA levels in rapamycin treated conditions relative to UT conditions is presented as the mean ± s.e.m. Expression of eIF4G1 and eIF4G2 mRNA was normalised to the housekeeping gene ALG9. (n = 3 independent biological replicates). n. s. (not significant). **(F)** Growth curves of WT cells overexpressing eIF4G1 or eIF4G2 on a plasmid grown in selective medium in the presence or absence of 200nM rapamycin. OD_600_ measurements were taken at 30-minute intervals for 24 hours.

In addition to suppressing translation initiation, TORC1 inhibition elicits a large transcriptional response. This is particularly well characterised for the ribosome biogenesis (RiBi) regulon, which consists of genes linked to ribosome biogenesis and translation (40, 41). The mRNA levels of these genes decrease in response to TORC1 inhibition, which, in addition to shut down of bulk translation, contributes to the reduction of their synthesis. eIF4G1 is known to be a constituent of the RiBi regulon, with its gene expression being rapidly downregulated following TORC1 inhibition. Similar regulation of eIF4G2 has not been established. To determine whether the defect in downregulating the protein levels of eIF4G in *zuo1Δ* cells is a result of transcriptional regulation, changes in the mRNA levels following rapamycin treatment were monitored using quantitative real-time PCR. As expected, gene expression of eIF4G1 is substantially downregulated upon rapamycin treatment in WT cells, whereas eIF4G2 mRNA levels appear to be largely unchanged by the treatment (Figure 7E). The transcriptional regulation of eIF4G1 and eIF4G2 does not seem to be impacted by the loss of Zuo1, suggesting that the defect in regulating eIF4G protein levels in *zuo1Δ* cells is occurring post-transcriptionally. This is consistent with the literature reporting that eIF4G proteins are selectively degraded upon TORC1 inhibition in yeast (42). To clarify whether the increased pool of eIF4G proteins observed in *zuo1Δ* cells treated with rapamycin is a cause or consequence of the proteostasis defect, the effect of overexpressing eIF4G1 and eIF4G2 in WT cells was examined. Over 24 hours, overexpression of either eIF4G paralog did not lead to a notable growth defect in WT cells under basal conditions (Figure 7F). On the other hand, overexpression of eIF4G causes a striking reduction of growth in WT cells grown in the presence of rapamycin, indicating that appropriate regulation of eIF4G is required for cells to survive rapamycin challenge. In contrast, the growth of *zuo1Δ* cells challenged with rapamycin is not further impaired by the overexpression of eIF4G-(Figure 7G). Altogether, these results suggest that failure to reduce eIF4G protein levels upon rapamycin treatment in *zuo1Δ* cells contributes to the loss of proteostasis.

## DISCUSSION

We have examined the role of Zuo1 in maintaining proteostasis in response to TORC1 inhibition. Cells lacking Zuo1 have a major defect in translational regulation, leading to a loss of proteostasis and cell death upon TORC1 inhibition. Shutting down bulk translation in response to TORC1 inhibition helps conserve resources when conditions are not optimal for growth, and it is therefore likely that loss of this regulation in *zuo1Δ* cells is causing the reduced viability in response to rapamycin treatment. Polysome levels in *zuo1Δ* cells are lower in basal conditions compared to WT cells, but the overall rate of translation was not perturbed, as previously reported (28, 29). Zuo1 has been linked to translation previously, where it is involved in translational repression of polylysine proteins (43) and, in mammalian cells, ribosome associated chaperones have been linked to elongation pausing upon stress (44, 45). It has also been proposed that this chaperone could slow down translation elongation to facilitate effective cotranslational protein folding (46, 47). It is therefore possible that, in the absence of Zuo1, translation elongation is proceeding faster than in WT cells, maintaining translation levels despite lower polysome levels. Failure to further decrease polysome levels upon challenge with rapamycin would therefore allow normal levels of translation to be sustained in *zuo1Δ* cells and disrupt its cellular proteostasis.

It appears that Zuo1 and Ssb cooperate to maintain proteostasis upon TORC1 inhibition. Surprisingly, though, the defect in reducing translation upon TORC1 inhibition is less severe in *zuo1Δssb1/2Δ* cells compared to *zuo1Δ* or *ssb1/2Δ* cells. Zuo1 and Ssb share binding sites on the ribosome with other ribosome associated chaperones, including NAC (35, 48, 49). It is possible that in the complete absence of RAC/Ssb, NAC is able to interact more with ribosomes and partially substitute for the loss of this chaperone system. It has been shown that NAC from *C. elegans* competes for ribosome binding with RAC (49) and, in yeast, NAC is expressed at approximately equimolar levels to ribosomes but is not bound to all translating ribosomes (50). In this context, the autonomous ribosomal binding of either Ssb or Zuo1 in *zuo1Δ* and *ssb1/2Δ* cells, respectively, may preclude the interaction of NAC, preventing a functional chaperone system from operating at those ribosomes. However, whether NAC shows increased association with translating ribosomes in *zuo1Δssb1/2Δ* cells remains to be determined. Additionally, deletion of NAC enhances growth defects of *ssb1/2Δ* cells (51), suggesting that these chaperones share overlapping functions. Further study is required to elucidate any interplay between these two chaperone systems for maintaining proteostasis upon TORC1 inhibition.

In the absence of TORC1 signalling, our results suggest that translation in *zuo1Δ* cells is partly sustained through elevated eIF4G levels. The argument for this is not unprecedented. It has previously been shown that mRNA decay mutants are able to sustain translation in response to a variety of stresses which induce translational shutdown (52). Although the mechanism has not been fully elucidated, it is thought that in these mutants, translation is maintained through increased eIF4F and mRNA interactions. In agreement with this, it was found that overexpression of eIF4G2 can prevent a decrease in polysomes, and active translation, in response to stress (52). Zuo1 has not been implicated in mRNA decay, but it is possible that loss of Zuo1 may similarly confer resistance to translational inhibition by facilitating ongoing translation through the eIF4F complex.

How eIF4G levels are maintained in *zuo1Δ* cells is currently unclear. It has been reported that eIF4G levels are decreased following TORC1 inhibition by increasing the rate of its degradation (42). Due to the positioning of Zuo1 on the ribosome and the observation that it interacts with eIF4G2 upon rapamycin treatment, it is attractive to speculate that Zuo1 is involved in promoting the turnover of eIF4G following TORC1 inhibition. One scenario is that Zuo1 could modulate post-translational modification of eIF4G which will serve as a recognition motif for degradation. We have not yet identified an interactor which could be involved in this process as none of the Zuo1 interactors analysed displayed both rapamycin sensitivity and proteostasis defect at the level of translation. Alternatively, Zuo1 and Ssb may have a more indirect role in promoting the folding or stability of an eIF4G modulator. Another scenario could involve Zuo1 regulating the production of de novo eIF4G. The nascent chains of both eIF4G paralogs have been identified as substrates of the RAC/Ssb chaperone system (15). Thus, upon TORC1 inhibition, Zuo1 may contribute to suppressing translation of eIF4G, preventing the pool of eIF4G from being replenished. Future work will focus on the elucidating the exact mechanism by which eIF4G levels remain elevated in *zuo1Δ* cells.

Inhibiting bulk translation is a key aspect of stress response under a large variety of stresses. This work has focused on TORC1 inhibition by the selective inhibitor rapamycin, but it would be useful to determine if Zuo1 plays a similar role in inhibiting translation following other stresses. This may indeed be the case as the deletion of Ssb prevents translation inhibition in response to glucose deprivation (53). As we have seen that Zuo1’s role relies on its ability to interact with Ssb, this chaperone complex may play a more ubiquitous role in regulating translation in response to stress.

Zuo1 is conserved in higher eukaryotes where it appears to share some functionality, as loss of this protein results in growth defects and hypersensitivity to aminoglycoside antibiotics, analogous to its loss in yeast (13, 54, 55). The mammalian homolog has also been implicated in regulating translation in basal conditions and has been linked to mTORC1 signalling (56, 57). It is therefore possible that the role of Zuo1 in regulating translation in response to TORC1 inhibition is not confined to yeast. Loss of proteostasis is linked to aging and disease progression, and so it will be important to understand the impact that the RAC complex plays in maintaining proteostasis under challenging conditions, including in cellular aging and disease contexts, in higher organisms.

## METHODS

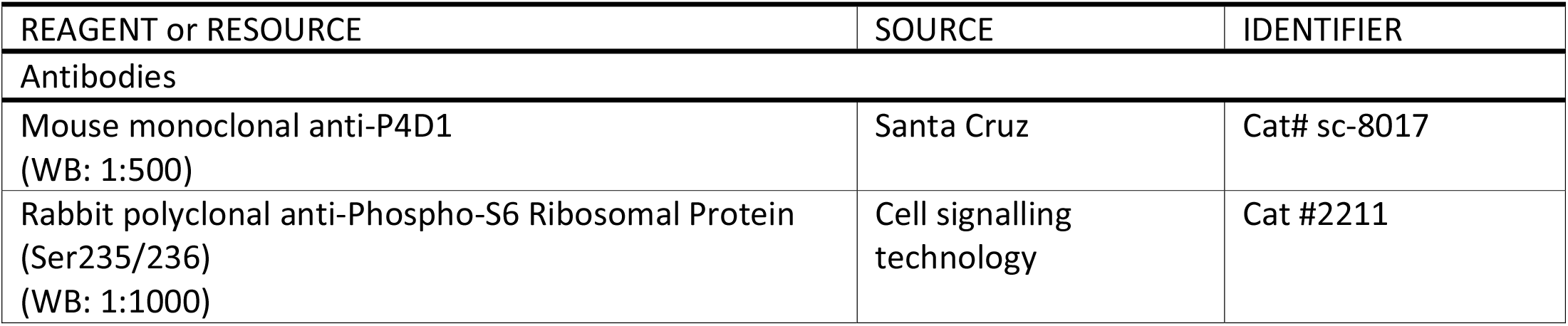
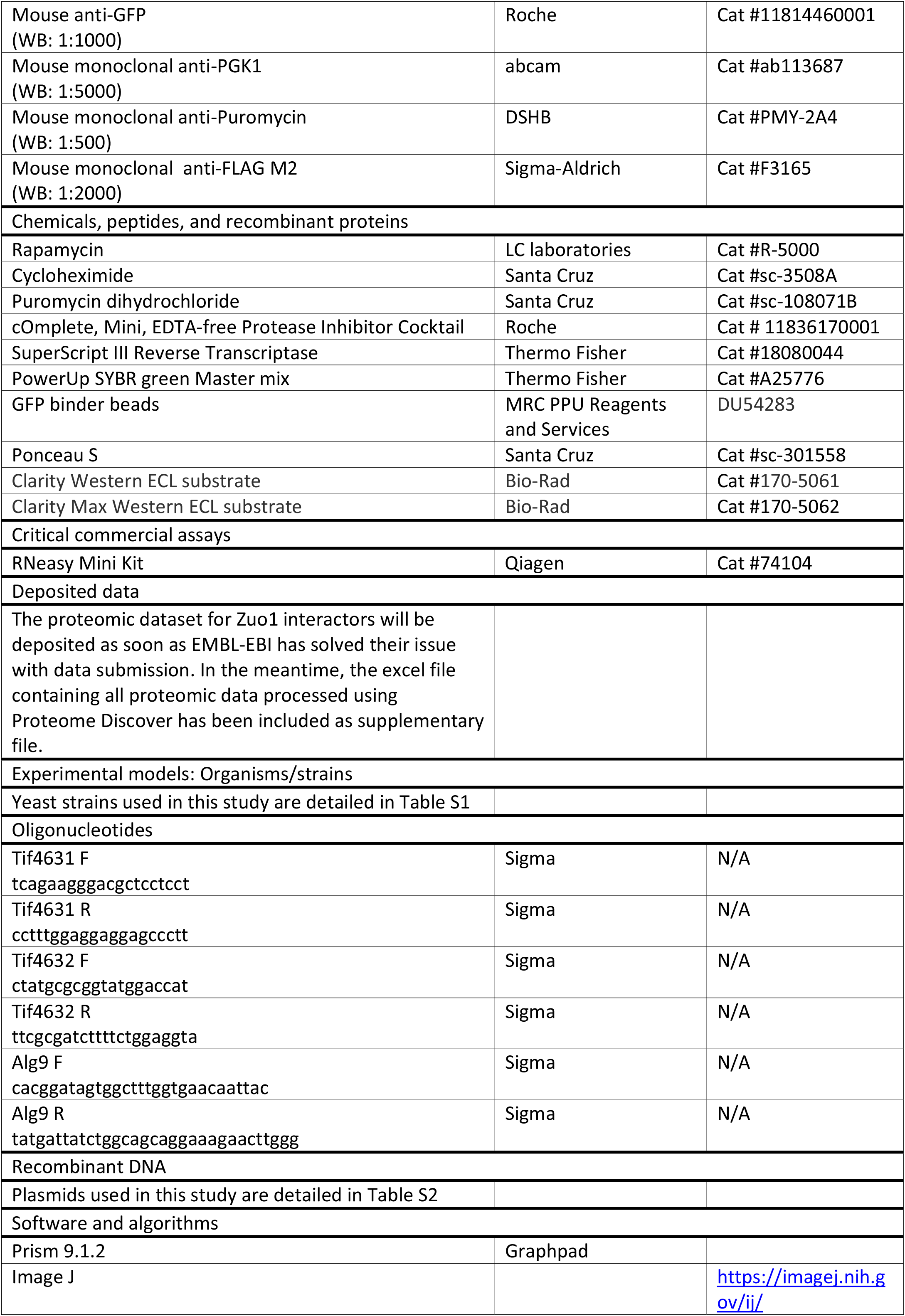
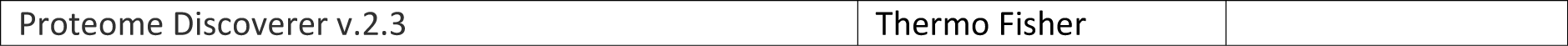

### Yeast strains, plasmids and growth conditions

All *Saccharomyces cerevisiae* strains used are isogenic to BY4741 and are listed in Table S1. Plasmids used are listed in Table S2 and were constructed using In-Fusion HD Cloning Plus. Gene deletions and genomic tagged strains were generated by homologous recombination as previously described (58). Genomic tagging with 5xFLAG was accomplished by PCR amplification of the 5xFLAG-His3MX cassette from pKL259. Integration of PCR cassettes was confirmed by PCR analysis. Yeast transformations were performed using the LiAc/PEG/SS carrier DNA method.

Yeast cells were cultured at 30°C in liquid medium with constant shaking at 200rpm. Cells were cultured overnight in YEPD (1% yeast extract, 2% peptone, 2% glucose) or synthetic medium (2% glucose) lacking the appropriate amino acids for plasmid selection and then readjusted to OD600 0.2. Cells were grown to mid-log phase and then readjusted to OD 0.2 prior to treatment with 200nM rapamycin or 37ug/ml cycloheximide. In the case of treatment with 0.5mM puromycin, yeast cells were readjusted to OD 0.3 instead of OD 0.2 on each occasion.

To assess growth cells, were equilibrated to OD_600_ 0.2 and either 5μl alone or 5-fold serial dilutions were spotted onto YEPD plates or YEPD plates containing 20ng/ml rapamycin and incubated at 30°C for 4 days. The Chemidoc MP imaging system (Bio-Rad) was used to image plates. To assess growth in liquid culture, cells were adjusted to OD 0.2 in YEPD and treated with rapamycin or vehicle control. Cells were cultured in triplicate in 98-well plates at 30°C and OD_600_ measurements were taken every 30 minutes over the course of 24 hours using the Fluostar Omega Microplate reader (BMG Labtech).

### Yeast protein extraction and western blot analysis

Cells were harvested (4000rpm, 4°C, 4 min), washed in ice-cold water (8000rpm, 4°C, 1 min), and flash frozen in dry ice then stored at −20°C before extraction. Yeast pellets were resuspended in 400μl 2M LiAC, on ice and centrifuged at 8000 rpm for 1 min. Pellets were then resuspended in 400μl 0.4M NaOH and centrifuged again before being resuspended in lysis buffer (0.1M NaOH, 0.05M EDTA, 2% SDS, 2% β-mercaptoethanol, and cOmplete protease inhibitor cocktail) and boiled at 90°C for 10 minutes. PhosStop was included in the lysis buffer for the detection of phosphorylated proteins, and N-ethylmaleimide was included for the detection of polyubiquitinated proteins. Acetic acid (4M) was then added (1:40), and samples were vortexed and boiled for an additional 10 minutes before centrifugation at 13,000 rpm for 15 minutes. The protein concentration of the supernatant was determined using a Nanodrop (A_280nm_, Thermo). The supernatant was combined with 5X loading buffer (0.25M Tris-HCl pH6.8, 50% glycerol, 0.05% bromophenol blue) and all samples were adjusted to the same protein concentration.

Approximately 30μg of each cell extract were separated on homemade 6-14% Bis-Tris acrylamide gradient gels (0.33M Bis-Tris pH 6.5) at 120V for 150 minutes at 4°C. Subsequently, gels were transferred onto 0.2μM nitrocellulose membranes (Bio-Rad) at 25V for 30 minutes using the Trans-blot Turbo transfer system (Bio-Rad). Ponceau S solution was used to stain membranes, which were imaged using the ChemiDoc MP system. Membranes were blocked in 5% (w/v) milk in TBS at room temperature for 1 hour, washed in TBS, and incubated in primary antibodies overnight at 4°C. Then, membranes were washed in TBS-T and incubated with secondary antibodies for 1 hour at room temperature prior to detection with Clarity ECL or Clarity Max ECL reagent using the ChemiDoc MP imaging system. Protein bands were quantified by densitometry using Image J, where indicated.

### In-gel peptidase assay

Exponential phase yeasts were adjusted to an OD_600_ 0.2 in 30ml and treated with 200nM rapamycin for 3h. Then, cells were harvested (400rpm, 4°C, 4 min) and washed in ice-cold water and centrifuged (8000rpm, 4°C, 1 min). The cell pellet was resuspended in native lysis buffer (50mM Tris pH8.5, 5mM MgCl_2_, 0.5mM EDTA, 5% glycerol, 1mM DTT, and 5mM ATP) and lysed with glass beads (FastPrep 24, 3×30s on/5min off). The supernatant was then cleared by centrifugation at 17,000g for 10 min at 4°C. Protein concentration was measured using a NanoDrop (A_280nm_, Thermo) and samples were adjusted to an equal concentration in native sample buffer (50mM Tris-HCl pH 6.8, 10% glycerol, 0.01% bromophenol blue). Samples were separated on 3.8-5% acrylamide native gel for 2h at 120V. Next gels were incubated with suc-LLVY-AMC fluorogenic substrate in assay buffer (50mM Tris-HCl pH7.5, 150mM NaCl, 5mM MgCl_2_, 10% glycerol) at 30°C for 20 minutes in the dark and subsequently imaged using the ChemiDoc MP system (Bio-Rad). The gel was then incubated in assay buffer for an additional 10 minutes with the addition of 0.1% SDS and imaged again.

### Polysome profiling

Polysome profile analysis was conducted as in (59) with some minor modifications. Yeast strains were cultured in YEPD medium overnight, diluted to OD_600_ 0.4 and grown until the OD_600_ reached 0.8-0.9. The cultures were then diluted to OD_600_ 0.4 and treated with 200nM rapamycin for 2h. 60OD of each culture was then taken and incubated with 100μg/ml cycloheximide on ice for 10 minutes. Cells were centrifuged at 4000 x g for 5 minutes at 4°C, washed in ice-cold buffer I (20mM Tris-HCl, pH7.4, 15mM MgCl_2_, 100mM KCl, 100μg/ml CHX) and centrifuged again. Next, the pellet was lysed in 250μl of buffer I supplemented with 1mM DTT, and EDTA free protease inhibitor cocktail using 250μl of acid washed glass beads and the FastPrep-24 (2×30 second cycles, 4m/sec, 4°C). Lysates were clarified by centrifugation at 17,000 x g for 15 minutes at 4°C. Seven A_260_ units were loaded onto 10-50% sucrose gradients, prepared using a Biocomp Gradient master instrument, in Buffer I containing 1mM DTT. The gradients were centrifuged at 35,000 rpm for 3 hours at 4°C (SW41 Ti rotor; Beckman) and subsequently fractionated using a Piston Gradient Fractionator (Biocomp instruments) monitoring absorbance at 260nm.

### Immunoprecipitation

Exponential phase yeasts were adjusted to an OD_600_ 0.2 in 30ml and treated with 200nM rapamycin for 3h. Then, cells were harvested (400rpm, 4°C, 4 min) and washed in ice-cold water and centrifuged (8000rpm, 4°C, 1 min). The cell pellet was resuspended in IP lysis buffer (50 mM Tris-HCl pH 8.0, 100 mM NaCl, 1 mM EDTA, 5 mM MgCl2, 1 mM DTT, 10% glycerol, 0.5 mM PMSF, and cOmplete protease inhibitor cocktail) and lysed with glass beads (FastPrep 24, 3×30s on/5min off). The supernatant was then cleared by centrifugation at 17,000g for 10 min at 4°C. Protein concentration was measured using a NanoDrop (A_280nm_, Thermo) and samples were equally adjusted to 1 mg/mL in IP lysis buffer. GFP binder beads (25 µl slurry per sample; MRC PPU reagents and services: DU54283) were equilibrated into 500 µl ice-cold wash buffer (10 mM Tris-HCl pH 7.5, 150 mM NaCl, and 0.5 mM EDTA) before being centrifuged at 2,500x g for 2 min at 4°C. Supernatant was discarded and equilibration steps repeated twice. Equilibrated beads were incubated with 1 mg of lysates for 1 h at 4 °C under rotation. GFP binder beads were then washed once with lysis buffer, twice with wash buffer. Beads were subjected to tryptic digestion and TMT-based quantitative proteomics (see below).

### On-bead tryptic digestion

Beads were collected and subsequently resuspended in 50 µl Urea buffer (2 M urea, 50 mM ammonium bicarbonate pH 8.0 and 5 mM DTT) and incubated at 45 °C for 30 min with gentle shaking in order to partially denature the proteins. Samples were centrifuged at 2,500g for 1 min and cooled to room temperature. Each sample was then incubated with iodoacetamide (10 mM final concentration) in the dark at room temperature. Unreacted iodoacetamide was then quenched with DTT (5 mM final concentration). Each sample was digested using 0.4 µg trypsin at 37 °C for 4 h under agitation. The digestion was stopped by adding trifluoroacetic acid (TFA) to the final 0.2% TFA concentration (v/v), centrifuged at 10,000g for 2 min at room temperature. The supernatant was de-salted on ultra-microspin column silica C18 (The Nest Group). De-salted peptides were dried using a SpeedVac vacuum centrifuge concentrator (Thermo Fisher) before TMT labelling.

### TMT labelling

Each vacuum-dried sample was resuspended in 50 μl of 100 mM TEAB buffer. The TMT labelling reagents were equilibrated to room temperature, and 41 μl anhydrous acetonitrile was added to each reagent channel and gently vortexed for 10 min. Then, 4 μl of each TMT reagent was added to the corresponding sample and labelling was performed at room temperature for 1 h with shaking before quenching with 1 μl of 5% hydroxylamine, after which 2 µl of labelled sample from each channel was analysed by liquid chromatography with tandem mass spectrometry (LC–MS/MS) to ensure complete labelling before mixing. After evaluation, the complete TMT-labelled samples were combined, acidified and dried. The mixture was then de-salted with ultra-microspin column silica C18, and the eluent from C18 column was dried.

### LC–MS/MS analysis

LC separations were performed with a Thermo Dionex Ultimate 3000 RSLC Nano liquid chromatography instrument. The dried peptides were dissolved in 0.1% formic acid and then loaded on C18 trap column with 3% ACN/0.1%TFA at a flow rate of 10 μl min−1. Peptide separations were performed using EASY-Spray columns (C18, 2 µm, 75 µm × 50 cm) with an integrated nano electrospray emitter at a flow rate of 300 nl min−1. Peptides were separated with a 180 min segmented gradient as follows starting from 8–30% buffer B in 135 min, ∼30–45% in 20 min and ∼45–90% in 5 min. Peptides eluted from the column were analysed on an Orbitrap Fusion Lumos (Thermo Fisher Scientific) mass spectrometer. Spray voltage was set to 2 kV, RF lens level was set at 30%, and ion transfer tube temperature was set to 275 °C. The Orbitrap Fusion Lumos was operated in positive-ion data-dependent mode with synchronous precursor selection (SPS)-MS3. The mass spectrometer was operated in data-dependent top speed mode with 3 s per cycle. The full scan was performed in the range of 350–1,500 m/z at nominal resolution of 120,000 at 200 m/z and AGC set to 4 × 105 with maximum injection time 50 ms, followed by selection of the most intense ions above an intensity threshold of 5 × 103 for collision-induced dissociation (CID)-MS2 fragmentation with 35% normalized collision energy. The isolation width was set to 0.7 m/z with no offset. Dynamic exclusion was set to 60 s. Monoisotopic precursor selection was set to peptide. Charge states between 2 and 7 were included for MS2 fragmentation. The MS2 scan was performed in the ion trap with auto normal range scan and AGC target of 1 × 104. The maximum injection time for MS2 scan was set to 50 ms. For the MS3 scan, SPS was enabled. MS3 was performed in the Orbitrap over 5 notches at a resolution of 50,000 at 200 m/z and AGC set to 5 × 104 with maximum injection time 105 ms, over a mass range of 100-500 m/z, with high collision induced dissociation (HCD) and 65% normalized collision energy.

### Proteomic data analysis

All the acquired LC–MS data were analysed using Proteome Discoverer v.2.3 (Thermo Fisher Scientific) with Mascot search engine. Maximum missed cleavage for trypsin digestion was set to 2. Precursor mass tolerance was set to 10 ppm. Fragment ion tolerance was set to 0.2 Da. Carbamidomethylation on cysteine (+57.021 Da) and TMT-10plex tags on N-termini as well as lysine (+229.163 Da) were set as static modifications. Variable modifications were set as oxidation on methionine (+15.995 Da). Data were searched against SwissProt database restricted to Saccharomyces cerevisiae taxonomy (reviewed 7905 entries downloaded in June 2019). Peptide spectral match error rates with a 1% false discovery rate were determined using the forward-decoy strategy modelling true and false matches.

Both unique and razor peptides were used for quantitation. Reporter ion abundances were corrected for isotopic impurities on the basis of the manufacturer’s data sheets. Reporter ions were quantified from MS3 scans using an integration tolerance of 20 ppm with the most confident centroid setting. Signal-to-noise (S/N) values were used to represent the reporter ion abundance with a co-isolation threshold of 50% and an average reporter S/N threshold of 10 and above required for quantitation from each MS3 spectra to be used. The summed abundance of quantified peptides was used for protein quantitation. The total peptide amount was used for the normalization. Protein ratios were calculated from medians of summed sample abundances of replicate groups. Standard deviation was calculated from all biological replicate values. The standard deviation of all biological replicates lower than 25% was used for further analyses. Multiple unpaired t-test was used to determine the significant differences between untreated and rapamycin-treated conditions.

### qRT-PCR

Total RNA from cells grown in the presence or absence of 200nM rapamycin for 2 hours was extracted using the RNeasy Kit according to the manufacturer’s instructions. Synthesis of cDNA from 100ng of total RNA was carried out using SuperScript III reverse transcriptase and random primers. qRT-PCR analysis was performed on a CFX384 real-time PCR detection system (Bio-Rad) using PowerUp SYBR Green Master Mix. Primer sequences are listed in the key resources table. The *ALG9* mRNA was used to normalise gene expression in each sample and changes in gene expression after rapamycin treatment were calculated using the ΔΔCt method.

### Statistical analysis

Each experiment was repeated independently a minimum of three times, as indicated. The standard deviation (s.d.) of the mean of at least three independent experiments is shown in the graphs, unless otherwise specified. *P* values are as stated, or not significant (NS). *P* values, numbers of independent biological replicates, and choice of statistical tests are described in individual figure legends. All statistics were performed using Graph Pad Prism 9 software (version 9.1.2) (Graph Pad Software Inc.). No statistical method was used to pre-determine sample size. No data were excluded from the analyses, and the experiments were not randomized.

## DATA AND CODE AVAILABILITY

The proteomic dataset for Zuo1 interactors will be deposited as soon as EMBL-EBI has solved their issue with data submission: “*EMBL-EBI storage system is facing some major issues affecting our submissions and publications jobs. The PRIDE team is working actively with EMBL-EBI storage team to solve these problems*”. In the meantime, the excel file containing processed proteomic data using Proteome Discoverer v.2.3 has been included as supplementary file.

This paper does not report original code.

## ACKNOWLEDGMENTS

We thank the MRC PPU reagents and services, including Cloning team and DNA sequencing services, MRC-PPU Mass Spectrometry and the Dundee Imaging facility. This research was supported by the Medical Research Council (grant number MC_UU_00018/8 to A.R.).

## AUTHOR CONTRIBUTIONS

Conceptualization, A.B., and A.R.; methodology, A.B., T.D.W., and A.R.; formal analysis, A.B., and A.R.; investigation, A.B., and A.R.; writing – original draft, A.B.; writing – review and editing, A.B., T.D.W., and A.R.; visualization, A.B., and A.R.; software, A.B., and A.R.; data curation, A.B., and A.R.; project administration, A.R.; supervision, A.R.; funding acquisition, A.R.

## INCLUSION AND DIVERSITY

We support inclusive, diverse, and equitable conduct of research.

---

**Figure S1.**
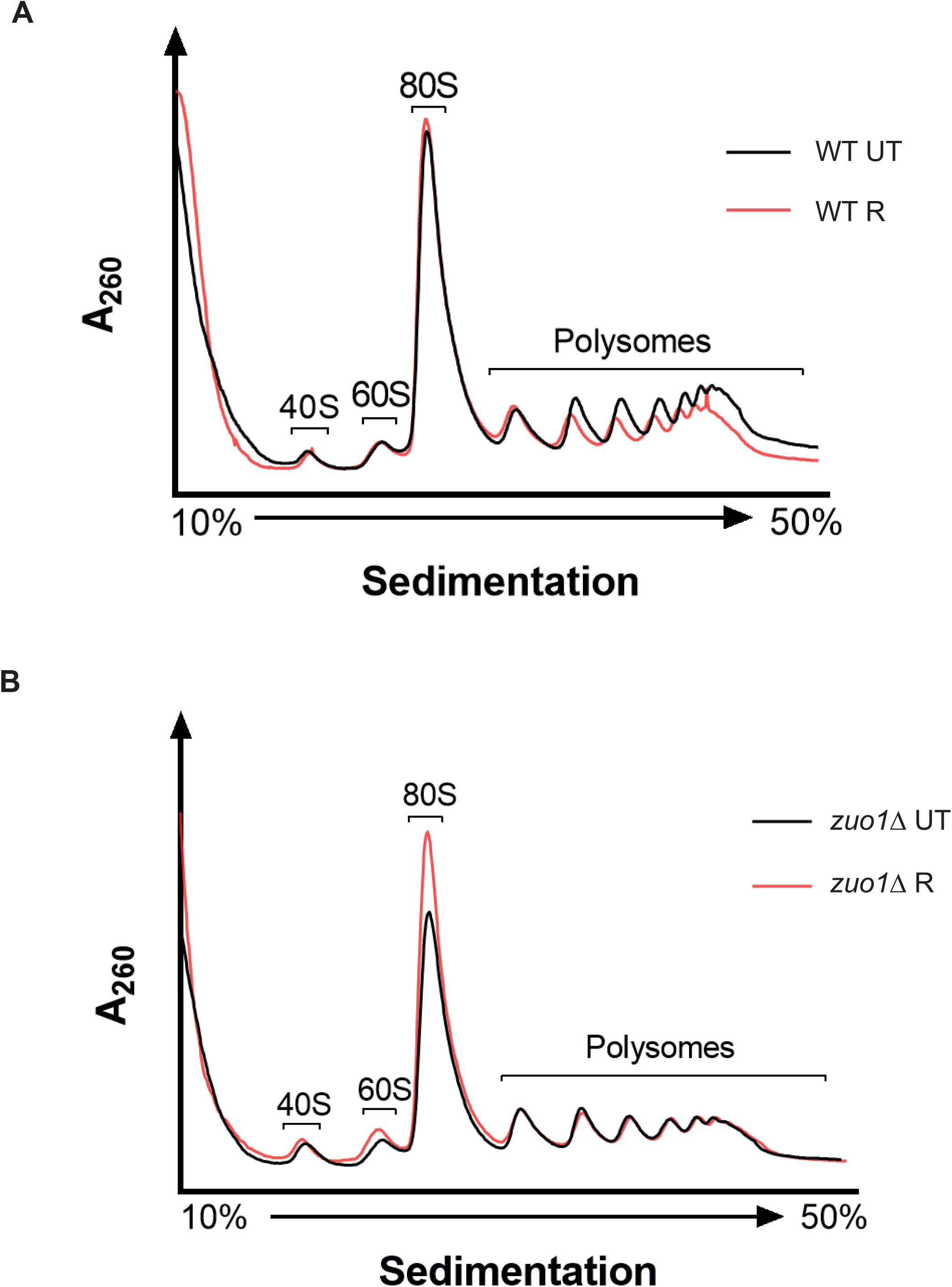
Polysome profiling analysis of WT and *zuo1*Δ cells. **(A) and (B)** Polysome profiles of WT (**A**) and *zuo1*Δ (**B**) strains either treated with 200nM rapamycin for 2 hours or left untreated. Total extracts were separated on 10-50% sucrose gradients and subsequently the A_260_ was monitored during fractionation.

**Figure S2.**
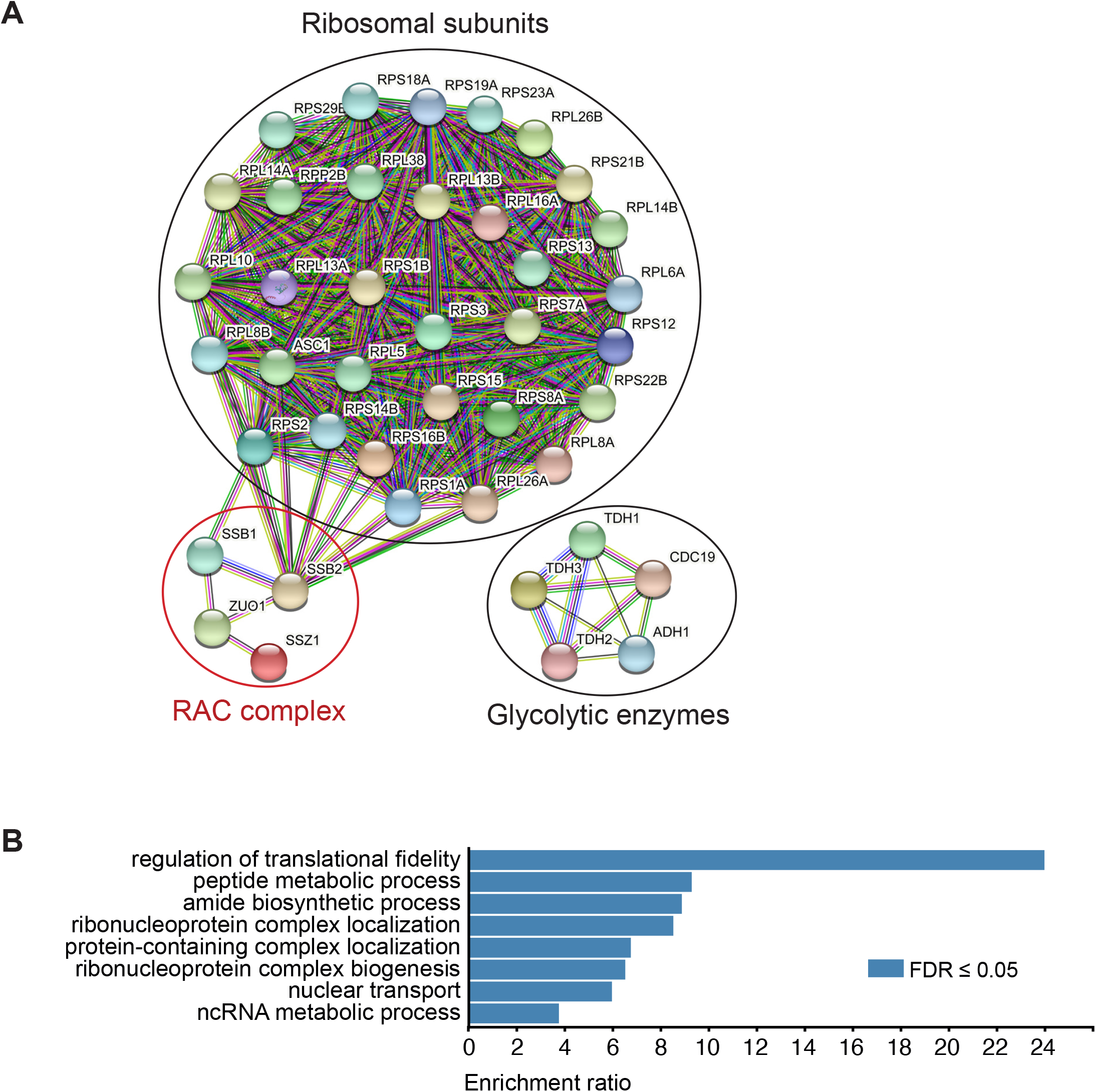
Gene ontology analysis of Zuo1 interactors. **(A)** Protein-protein interaction network of Zuo1 interactors having at least 50% peptide coverage. **(B)** Gene ontology analysis of Zuo1 interactors having at least 50% peptide coverage.

**Figure S3.**
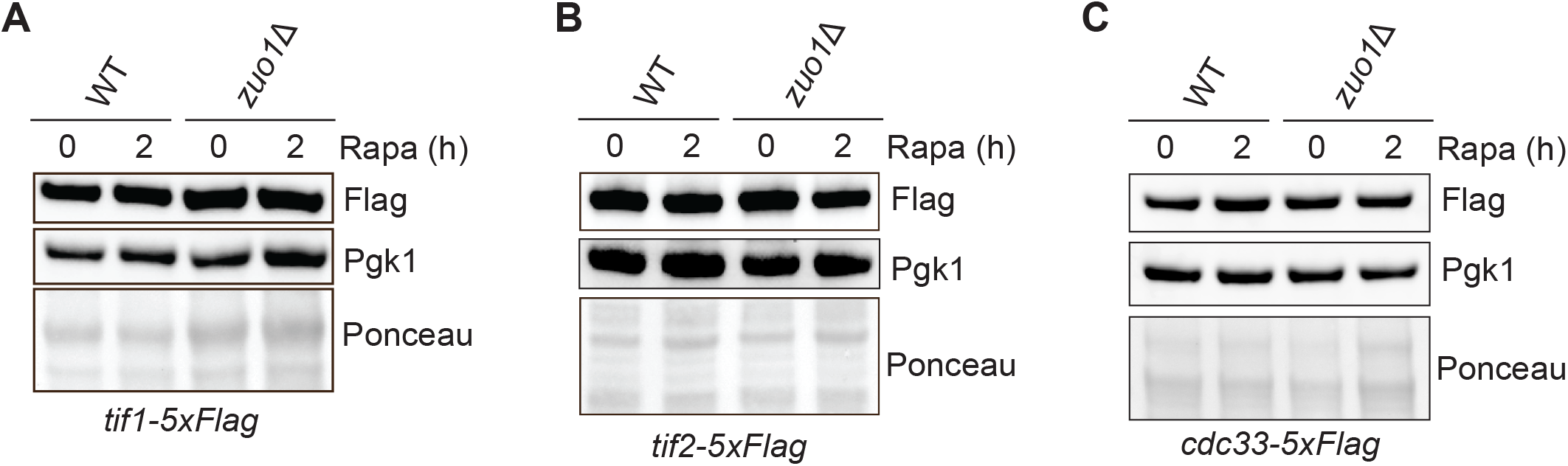
Abundance of eIF4F subunits upon TORC1 inhibition. **(A)**, **(B) and (C)** Immunoblot analysis of lysates from WT and *zuo1*Δ cells containing TIF1-5xFLAG (**A**), TIF2-5xFLAG (**B**), or CDC33-5xFLAG (**C**) at the endogenous locus treated with 200nM rapamycin for 2 hours or left untreated. Ponceau and Pgk1 staining served as the loading control.

**Supplementary Table 1.**
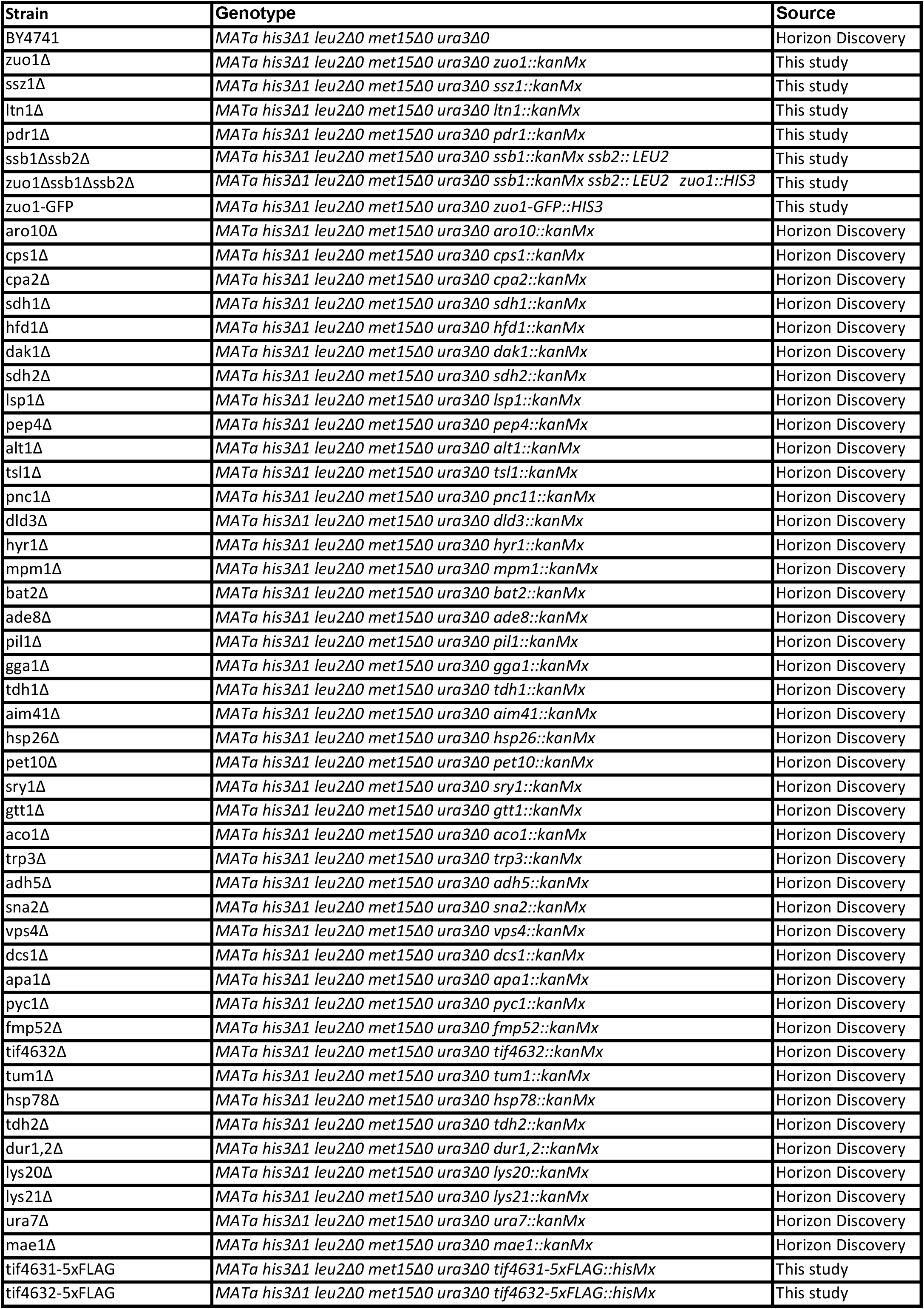
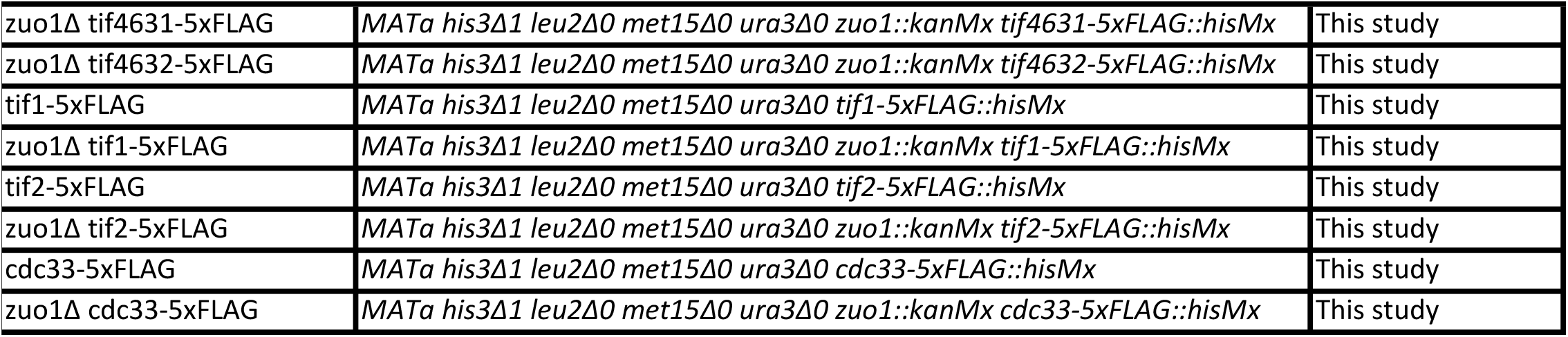
List of strains used in this study.

| Strain | Genotype | Source |
| --- | --- | --- |
| BY4741 | *MATa his3Δ1 leu2Δ0 met15Δ0 ura3Δ0* | Horizon Discovery |
| zuo1Δ | *MATa his3Δ1 leu2Δ0 met15Δ0 ura3Δ0 zuo1::kanMx* | This study |
| ssz1Δ | *MATa his3Δ1 leu2Δ0 met15Δ0 ura3Δ0 ssz1::kanMx* | This study |
| ltn1Δ | *MATa his3Δ1 leu2Δ0 met15Δ0 ura3Δ0 ltn1::kanMx* | This study |
| pdr1Δ | *MATa his3Δ1 leu2Δ0 met15Δ0 ura3Δ0 pdr1::kanMx* | This study |
| ssb1Δssb2Δ | *MATa his3Δ1 leu2Δ0 met15Δ0 ura3Δ0 ssb1::kanMx ssb2::LEU2* | This study |
| zuo1Δssb1Δssb2Δ | *MATa his3Δ1 leu2Δ0 met15Δ0 ura3Δ0 ssb1::kanMx ssb2::LEU2 zuo1::HIS3* | This study |
| zuo1-GFP | *MATa his3Δ1 leu2Δ0 met15Δ0 ura3Δ0 zuo1-GFP::HIS3* | This study |
| aro10Δ | *MATa his3Δ1 leu2Δ0 met15Δ0 ura3Δ0 aro10::kanMx* | Horizon Discovery |
| cps1Δ | *MATa his3Δ1 leu2Δ0 met15Δ0 ura3Δ0 cps1::kanMx* | Horizon Discovery |
| cpa2Δ | *MATa his3Δ1 leu2Δ0 met15Δ0 ura3Δ0 cpa2::kanMx* | Horizon Discovery |
| sdh1Δ | *MATa his3Δ1 leu2Δ0 met15Δ0 ura3Δ0 sdh1::kanMx* | Horizon Discovery |
| hfd1Δ | *MATa his3Δ1 leu2Δ0 met15Δ0 ura3Δ0 hfd1::kanMx* | Horizon Discovery |
| dak1Δ | *MATa his3Δ1 leu2Δ0 met15Δ0 ura3Δ0 dak1::kanMx* | Horizon Discovery |
| sdh2Δ | *MATa his3Δ1 leu2Δ0 met15Δ0 ura3Δ0 sdh2::kanMx* | Horizon Discovery |
| lsp1Δ | *MATa his3Δ1 leu2Δ0 met15Δ0 ura3Δ0 lsp1::kanMx* | Horizon Discovery |
| pep4Δ | *MATa his3Δ1 leu2Δ0 met15Δ0 ura3Δ0 pep4::kanMx* | Horizon Discovery |
| alt1Δ | *MATa his3Δ1 leu2Δ0 met15Δ0 ura3Δ0 alt1::kanMx* | Horizon Discovery |
| tsl1Δ | *MATa his3Δ1 leu2Δ0 met15Δ0 ura3Δ0 tsl1::kanMx* | Horizon Discovery |
| pnc1Δ | *MATa his3Δ1 leu2Δ0 met15Δ0 ura3Δ0 pnc1::kanMx* | Horizon Discovery |
| dld3Δ | *MATa his3Δ1 leu2Δ0 met15Δ0 ura3Δ0 dld3::kanMx* | Horizon Discovery |
| hyr1Δ | *MATa his3Δ1 leu2Δ0 met15Δ0 ura3Δ0 hyr1::kanMx* | Horizon Discovery |
| mpm1Δ | *MATa his3Δ1 leu2Δ0 met15Δ0 ura3Δ0 mpm1::kanMx* | Horizon Discovery |
| bat2Δ | *MATa his3Δ1 leu2Δ0 met15Δ0 ura3Δ0 bat2::kanMx* | Horizon Discovery |
| ade8Δ | *MATa his3Δ1 leu2Δ0 met15Δ0 ura3Δ0 ade8::kanMx* | Horizon Discovery |
| pil1Δ | *MATa his3Δ1 leu2Δ0 met15Δ0 ura3Δ0 pil1::kanMx* | Horizon Discovery |
| gga1Δ | *MATa his3Δ1 leu2Δ0 met15Δ0 ura3Δ0 gga1::kanMx* | Horizon Discovery |
| tdh1Δ | *MATa his3Δ1 leu2Δ0 met15Δ0 ura3Δ0 tdh1::kanMx* | Horizon Discovery |
| aim41Δ | *MATa his3Δ1 leu2Δ0 met15Δ0 ura3Δ0 aim41::kanMx* | Horizon Discovery |
| hsp26Δ | *MATa his3Δ1 leu2Δ0 met15Δ0 ura3Δ0 hsp26::kanMx* | Horizon Discovery |
| pet10Δ | *MATa his3Δ1 leu2Δ0 met15Δ0 ura3Δ0 pet10::kanMx* | Horizon Discovery |
| sry1Δ | *MATa his3Δ1 leu2Δ0 met15Δ0 ura3Δ0 sry1::kanMx* | Horizon Discovery |
| gtt1Δ | *MATa his3Δ1 leu2Δ0 met15Δ0 ura3Δ0 gtt1::kanMx* | Horizon Discovery |
| aco1Δ | *MATa his3Δ1 leu2Δ0 met15Δ0 ura3Δ0 aco1::kanMx* | Horizon Discovery |
| trp3Δ | *MATa his3Δ1 leu2Δ0 met15Δ0 ura3Δ0 trp3::kanMx* | Horizon Discovery |
| adh5Δ | *MATa his3Δ1 leu2Δ0 met15Δ0 ura3Δ0 adh5::kanMx* | Horizon Discovery |
| sna2Δ | *MATa his3Δ1 leu2Δ0 met15Δ0 ura3Δ0 sna2::kanMx* | Horizon Discovery |
| vps4Δ | *MATa his3Δ1 leu2Δ0 met15Δ0 ura3Δ0 vps4::kanMx* | Horizon Discovery |
| dcs1Δ | *MATa his3Δ1 leu2Δ0 met15Δ0 ura3Δ0 dcs1::kanMx* | Horizon Discovery |
| apa1Δ | *MATa his3Δ1 leu2Δ0 met15Δ0 ura3Δ0 apa1::kanMx* | Horizon Discovery |
| pyc1Δ | *MATa his3Δ1 leu2Δ0 met15Δ0 ura3Δ0 pyc1::kanMx* | Horizon Discovery |
| fmp52Δ | *MATa his3Δ1 leu2Δ0 met15Δ0 ura3Δ0 fmp52::kanMx* | Horizon Discovery |
| tif4632Δ | *MATa his3Δ1 leu2Δ0 met15Δ0 ura3Δ0 tif4632::kanMx* | Horizon Discovery |
| tum1Δ | *MATa his3Δ1 leu2Δ0 met15Δ0 ura3Δ0 tum1::kanMx* | Horizon Discovery |
| hsp78Δ | *MATa his3Δ1 leu2Δ0 met15Δ0 ura3Δ0 hsp78::kanMx* | Horizon Discovery |
| tdh2Δ | *MATa his3Δ1 leu2Δ0 met15Δ0 ura3Δ0 tdh2::kanMx* | Horizon Discovery |
| dur1,2Δ | *MATa his3Δ1 leu2Δ0 met15Δ0 ura3Δ0 dur1,2::kanMx* | Horizon Discovery |
| lys20Δ | *MATa his3Δ1 leu2Δ0 met15Δ0 ura3Δ0 lys20::kanMx* | Horizon Discovery |
| lys21Δ | *MATa his3Δ1 leu2Δ0 met15Δ0 ura3Δ0 lys21::kanMx* | Horizon Discovery |
| ura7Δ | *MATa his3Δ1 leu2Δ0 met15Δ0 ura3Δ0 ura7::kanMx* | Horizon Discovery |
| mae1Δ | *MATa his3Δ1 leu2Δ0 met15Δ0 ura3Δ0 mae1::kanMx* | Horizon Discovery |
| tif4631-5xFLAG | *MATa his3Δ1 leu2Δ0 met15Δ0 ura3Δ0 tif4631-5xFLAG::hisMx* | This study |
| tif4632-5xFLAG | *MATa his3Δ1 leu2Δ0 met15Δ0 ura3Δ0 tif4632-5xFLAG::hisMx* | This study |
| zuo1Δ tif4631-5xFLAG | *MATa his3Δ1 leu2Δ0 met15Δ0 ura3Δ0 zuo1::kanMx tif4631-5xFLAG::hisMx* | This study |
| zuo1Δ tif4632-5xFLAG | *MATa his3Δ1 leu2Δ0 met15Δ0 ura3Δ0 zuo1::kanMx tif4632-5xFLAG::hisMx* | This study |
| tif1-5xFLAG | *MATa his3Δ1 leu2Δ0 met15Δ0 ura3Δ0 tif1-5xFLAG::hisMx* | This study |
| zuo1Δ tif1-5xFLAG | *MATa his3Δ1 leu2Δ0 met15Δ0 ura3Δ0 zuo1::kanMx tif1-5xFLAG::hisMx* | This study |
| tif2-5xFLAG | *MATa his3Δ1 leu2Δ0 met15Δ0 ura3Δ0 tif2-5xFLAG::hisMx* | This study |
| zuo1Δ tif2-5xFLAG | *MATa his3Δ1 leu2Δ0 met15Δ0 ura3Δ0 zuo1::kanMx tif2-5xFLAG::hisMx* | This study |
| cdc33-5xFLAG | *MATa his3Δ1 leu2Δ0 met15Δ0 ura3Δ0 cdc33-5xFLAG::hisMx* | This study |
| zuo1Δ cdc33-5xFLAG | *MATa his3Δ1 leu2Δ0 met15Δ0 ura3Δ0 zuo1::kanMx cdc33-5xFLAG::hisMx* | This study |

**Supplementary Table 2.** List of plasmids used in this study.

| Insert | Vector backbone | Source |
| --- | --- | --- |
| Δss-CPY*GFP | p416-GPD | This Study |
|  | p416-GPD |  |
| ZUO1 | p416-GPD | This Study |
| ZUO1-R247A/R251A | p416-GPD | This Study |
| ZUO1-H128A/P129A/D130A | p416-GPD | This Study |
| ZUO1-R247A/R251A/<br>H128A/P129A/D130A | p416-GPD | This Study |
| TIF4631 | p416-GPD | This Study |
| TIF4632 | p416-GPD | This Study |

## REFERENCES

1. Labbadia J, Morimoto RI. 2015. The Biology of Proteostasis in Aging and Disease. Annu Rev Biochem 84:435.

2. Klaips CL, Jayaraj GG, Hartl FU. 2018. Pathways of cellular proteostasis in aging and disease. Journal of Cell Biology 217:51–63.

3. Ben-Sahra I, Manning BD. 2017. mTORC1 signaling and the metabolic control of cell growth. Curr Opin Cell Biol 45:72.

4. The TOR signaling cascade regulates gene expression in response to nutrients. http://genesdev.cshlp.org/content/13/24/3271. Retrieved 17 August 2022.

5. Lippman SI, Broach JR. 2009. Protein kinase A and TORC1 activate genes for ribosomal biogenesis by inactivating repressors encoded by Dot6 and its homolog Tod6. Proc Natl Acad Sci U S A 106:19928.

6. Powers T, Walter P. 1999. Regulation of Ribosome Biogenesis by the Rapamycin-sensitive TOR-signaling Pathway in Saccharomyces cerevisiae. Mol Biol Cell 10:987.

7. Kamada Y, Yoshino K, Kondo C, Kawamata T, Oshiro N, Yonezawa K, Ohsumi Y. 2010. Tor Directly Controls the Atg1 Kinase Complex To Regulate Autophagy. Mol Cell Biol 30:1049–1058.

8. Rousseau A, Bertolotti A. 2016. An evolutionarily conserved pathway controls proteasome homeostasis. Nature 536:184–189.

9. Williams TD, Cacioppo R, Agrotis A, Black A, Zhou H, Rousseau A. 2022. Actin remodelling controls proteasome homeostasis upon stress. Nature Cell Biology 2022 24:7 24:1077–1087.

10. Hartl FU, Bracher A, Hayer-Hartl M. 2011. Molecular chaperones in protein folding and proteostasis. Nature 2011 475:7356 475:324–332.

11. Gautschi M, Mun A, Ross S, Rospert S. 2002. A functional chaperone triad on the yeast ribosome. Proc Natl Acad Sci U S A 99:4209–4214.

12. Zhang Y, Sinning I, Rospert S. 2017. Two chaperones locked in an embrace: structure and function of the ribosome-associated complex RAC. Nature Structural & Molecular Biology 2017 24:8 24:611–619.

13. Jaiswal H, Conz C, Otto H, Wölfle T, Fitzke E, Mayer MP, Rospert S. 2011. The Chaperone Network Connected to Human Ribosome-Associated Complex. Mol Cell Biol 31:1160.

14. Willmund F, del Alamo M, Pechmann S, Chen T, Albanèse V, Dammer EB, Peng J, Frydman J. 2013. The cotranslational function of ribosome-associated Hsp70 in eukaryotic protein homeostasis. Cell 152:196.

15. Döring K, Ahmed N, Riemer T, Suresh HG, Vainshtein Y, Habich M, Riemer J, Mayer MP, O’Brien EP, Kramer G, Bukau B. 2017. Profiling Ssb-nascent chain interactions reveals principles of Hsp70 assisted folding. Cell 170:298.

16. Kolkman A, Daran-Lapujade P, Fullaondo A, Olsthoorn MMA, Pronk JT, Slijper M, Heck AJR. 2006. Proteome analysis of yeast response to various nutrient limitations. Mol Syst Biol 2.

17. Shenton D, Smirnova JB, Selley JN, Carroll K, Hubbard SJ, Pavitt GD, Ashe MP, Grant CM. 2006. Global translational responses to oxidative stress impact upon multiple levels of protein synthesis. J Biol Chem 281:29011–29021.

18. Causton HC, Ren B, Sang Seok Koh, Harbison CT, Kanin E, Jennings EG, Tong Ihn Lee, True HL, Lander ES, Young RA. 2001. Remodeling of yeast genome expression in response to environmental changes. Mol Biol Cell 12:323–337.

19. Simpson CE, Ashe MP. 2012. Adaptation to stress in yeast: to translate or not? Biochem Soc Trans 40:794–799.

20. Melamed D, Pnueli L, Arava Y. 2008. Yeast translational response to high salinity: Global analysis reveals regulation at multiple levels. RNA 14:1337.

21. Ashe MP, de Long SK, Sachs AB. 2000. Glucose Depletion Rapidly Inhibits Translation Initiation in Yeast. Mol Biol Cell 11:833.

22. Preiss T, Baron-Benhamou J, Ansorge W, Hentze MW. 2003. Homodirectional changes in transcriptome composition and mRNA translation induced by rapamycin and heat shock. Nature Structural & Molecular Biology 2003 10:12 10:1039–1047.

23. Gillies AT, Taylor R, Gestwicki JE. 2012. Synthetic lethal interactions in yeast reveal functional roles of J protein co-chaperones. Mol Biosyst 8:2901.

24. Yerlikaya S, Meusburger M, Kumari R, Huber A, Anrather D, Costanzo M, Boone C, Ammerer G, Baranov P v., Loewith R. 2016. TORC1 and TORC2 work together to regulate ribosomal protein S6 phosphorylation in Saccharomyces cerevisiae. Mol Biol Cell 27:397.

25. Medicherla B, Kostova Z, Schaefer A, Wolf DH. 2004. A genomic screen identifies Dsk2p and Rad23p as essential components of ER-associated degradation. EMBO Rep 5:692.

26. Waite KA, Burris A, Vontz G, Lang A, Roelofs J. 2022. Proteaphagy is specifically regulated and requires factors dispensable for general autophagy. J Biol Chem 298.

27. Schmidt EK, Clavarino G, Ceppi M, Pierre P. 2009. SUnSET, a nonradioactive method to monitor protein synthesis. Nat Methods 6:275–277.

28. Albanèse V, Reissmann S, Frydman J. 2010. A ribosome-anchored chaperone network that facilitates eukaryotic ribosome biogenesis. Journal of Cell Biology 189:69–81.

29. Hanebuth MA, Kityk R, Fries SJ, Jain A, Kriel A, Albanese V, Frickey T, Peter C, Mayer MP, Frydman J, Deuerling E. 2016. Multivalent contacts of the Hsp70 Ssb contribute to its architecture on ribosomes and nascent chain interaction. Nature Communications 2016 7:1 7:1–13.

30. Ghosh A, Shcherbik N. 2020. Cooperativity between the Ribosome-Associated Chaperone Ssb/RAC and the Ubiquitin Ligase Ltn1 in Ubiquitination of Nascent Polypeptides. Int J Mol Sci 21:1–22.

31. Eisenman HC, Craig EA. 2004. Activation of pleiotropic drug resistance by the J-protein and Hsp70-related proteins, Zuo1 and Ssz1. Mol Microbiol 53:335–344.

32. Hundley H, Eisenman H, Walter W, Evans T, Hotokezaka Y, Wiedmann M, Craig E. 2002. The in vivo function of the ribosome-associated Hsp70, Ssz1, does not require its putative peptide-binding domain. Proc Natl Acad Sci U S A 99:4203–4208.

33. Kaschner LA, Sharma R, Shrestha OK, Meyer AE, Craig EA. 2015. A conserved domain important for association of eukaryotic J-protein co-chaperones Jjj1 and Zuo1 with the ribosome. Biochim Biophys Acta 1853:1035.

34. Huang P, Gautschi M, Walter W, Rospert S, Craig EA. 2005. The Hsp70 Ssz1 modulates the function of the ribosome-associated J-protein Zuo1. Nature Structural & Molecular Biology 2005 12:6 12:497–504.

35. Gumiero A, Conz C, Gesé GV, Zhang Y, Weyer FA, Lapouge K, Kappes J, von Plehwe U, Schermann G, Fitzke E, Wölfle T, Fischer T, Rospert S, Sinning I. 2016. Interaction of the cotranslational Hsp70 Ssb with ribosomal proteins and rRNA depends on its lid domain. Nature Communications 2016 7:1 7:1–12.

36. Hinnebusch AG, Lorsch JR. 2012. The Mechanism of Eukaryotic Translation Initiation: New Insights and Challenges. Cold Spring Harb Perspect Biol 4.

37. Berset C, Trachsel H, Altmann M. 1998. The TOR (target of rapamycin) signal transduction pathway regulates the stability of translation initiation factor eIF4G in the yeast Saccharomyces cerevisiae. Proc Natl Acad Sci U S A 95:4264.

38. Gilbert W v., Zhou K, Butler TK, Doudna JA. 2007. Cap-independent translation is required for starvation-induced differentiation in yeast. Science 317:1224–1227.

39. von der Haar T, McCarthy JEG. 2002. Intracellular translation initiation factor levels in Saccharomyces cerevisiae and their role in cap-complex function. Mol Microbiol 46:531–544.

40. Hardwick JS, Kuruvilla FG, Tong JK, Shamji AF, Schreiber SL. 1999. Rapamycin-modulated transcription defines the subset of nutrient-sensitive signaling pathways directly controlled by the Tor proteins. Proc Natl Acad Sci U S A 96:14866–14870.

41. Powers T, Walter P. 1999. Regulation of ribosome biogenesis by the rapamycin-sensitive TOR-signaling pathway in Saccharomyces cerevisiae. Mol Biol Cell 10:987–1000.

42. Kelly SP, Bedwell DM. 2015. Both the autophagy and proteasomal pathways facilitate the Ubp3p-dependent depletion of a subset of translation and RNA turnover factors during nitrogen starvation in Saccharomyces cerevisiae. RNA 21:898.

43. Chiabudini M, Conz C, Reckmann F, Rospert S. 2012. Ribosome-Associated Complex and Ssb Are Required for Translational Repression Induced by Polylysine Segments within Nascent Chains. Mol Cell Biol 32:4769.

44. Shalgi R, Hurt JA, Krykbaeva I, Taipale M, Lindquist S, Burge CB. 2013. Widespread Regulation of Translation by Elongation Pausing in Heat Shock. Mol Cell 49:439–452.

45. Liu B, Han Y, Qian SB. 2013. Cotranslational Response to Proteotoxic Stress by Elongation Pausing of Ribosomes. Mol Cell 49:453–463.

46. Zhang Y, Ma C, Yuan Y, Zhu J, Li N, Chen C, Wu S, Yu L, Lei J, Gao N. 2014. Structural basis for interaction of a cotranslational chaperone with the eukaryotic ribosome. Nature Structural & Molecular Biology 2014 21:12 21:1042–1046.

47. Chen Y, Tsai B, Li N, Gao N. 2022. Structural remodeling of ribosome associated Hsp40-Hsp70 chaperones during co-translational folding. Nature Communications 2022 13:1 13:1–14.

48. Pech M, Spreter T, Beckmann R, Beatrix B. 2010. Dual Binding Mode of the Nascent Polypeptide-associated Complex Reveals a Novel Universal Adapter Site on the Ribosome. Journal of Biological Chemistry 285:19679–19687.

49. Gamerdinger M, Kobayashi K, Wallisch A, Kreft SG, Sailer C, Schlömer R, Sachs N, Jomaa A, Stengel F, Ban N, Deuerling E. 2019. Early Scanning of Nascent Polypeptides inside the Ribosomal Tunnel by NAC. Mol Cell 75:996-1006.e8.

50. Raue U, Oellerer S, Rospert S. 2007. Association of Protein Biogenesis Factors at the Yeast Ribosomal Tunnel Exit Is Affected by the Translational Status and Nascent Polypeptide Sequence. Journal of Biological Chemistry 282:7809–7816.

51. Koplin, Ansgar. 2010. A dual function for chaperones SSB–RAC and the NAC nascent polypeptide–associated complex on ribosomes https://doi.org/10.1083/jcb.200910074.

52. Holmes LEA, Campbell SG, Long SK de, Sachs AB, Ashe MP. 2004. Loss of Translational Control in Yeast Compromised for the Major mRNA Decay Pathway. Mol Cell Biol 24:2998.

53. von Plehwe U, Berndt U, Conz C, Chiabudini M, Fitzke E, Sickmann A, Petersen A, Pfeifer D, Rospert S. 2009. The Hsp70 homolog Ssb is essential for glucose sensing via the SNF1 kinase network. Genes Dev 23:2102.

54. Aloia L, di Stefano B, Sessa A, Morey L, Santanach A, Gutierrez A, Cozzuto L, Benitah SA, Graf T, Broccoli V, di Croce L. 2014. Zrf1 is required to establish and maintain neural progenitor identity. Genes Dev 28:182–197.

55. Shoji W, Inoue T, Yamamoto T, Obinata M. 1995. MIDA1, a protein associated with Id, regulates cell growth. Journal of Biological Chemistry 270:24818–24825.

56. Barilari M, Bonfils G, Treins C, Koka V, Villeneuve D de, Fabrega S, Pende M. 2017. ZRF1 is a novel S6 kinase substrate that drives the senescence programme. EMBO J 36:736–750.

57. Wu IH, Yoon JS, Yang Q, Liu Y, Skach W, Thomas P. 2021. A role for the ribosome-associated complex in activation of the IRE1 branch of UPR. Cell Rep 35:109217.

58. Janke C, Magiera MM, Rathfelder N, Taxis C, Reber S, Maekawa H, Moreno-Borchart A, Doenges G, Schwob E, Schiebel E, Knop M. 2004. A versatile toolbox for PCR-based tagging of yeast genes: new fluorescent proteins, more markers and promoter substitution cassettes. Yeast 21:947–962.

59. Kasari V, Margus T, Atkinson GC, Johansson MJO, Hauryliuk V. 2019. Ribosome profiling analysis of eEF3-depleted Saccharomyces cerevisiae. Scientific Reports 2019 9:1 9:1–10.

